# Human NOP2/NSUN1 Regulates Ribosome Biogenesis Through Non-Catalytic Complex Formation with Box C/D snoRNPs

**DOI:** 10.1101/2021.11.12.468419

**Authors:** Han Liao, Anushri Gaur, Hunter McConie, Amirtha Shekar, Karen Wang, Jeffrey T. Chang, Ghislain Breton, Catherine Denicourt

## Abstract

5-Methylcytosine (m^5^C) is a base modification broadly found on various RNAs in the human transcriptome. In eukaryotes, m^5^C is catalyzed by enzymes of the NSUN family composed of seven human members (NSUN1-7). NOP2/NSUN1 has been primarily characterized in budding yeast as an essential ribosome biogenesis factor required for the deposition of m^5^C on the 25S ribosomal RNA (rRNA). Although human NOP2/NSUN1 has been known to be an oncogene overexpressed in several types of cancer, its functions and substrates remain poorly characterized. Here we used a miCLIP-seq approach to identify human NOP2/NSUN1 RNA substrates. Our analysis revealed that NOP2/NSUN1 catalyzes the deposition of m^5^C at position 4447 on the 28S rRNA. We also find that NOP2/NSUN1 binds to the 5’ETS region of the pre-rRNA transcript and regulates pre-rRNA processing through non-catalytic complex formation with box C/D snoRNAs. We provide evidence that NOP2/NSUN1 facilitates the recruitment of U3 and U8 snoRNAs to pre-90S ribosomal particles and their stable assembly into snoRNP complexes. Remarkably, expression of both WT and catalytically inactive NOP2/NSUN1 in knockdown background rescues the rRNA processing defects and the stable assembly of box C/D snoRNP complexes, suggesting that NOP2/NSUN1-mediated deposition of m^5^C on rRNA is not required for ribosome synthesis.

## INTRODUCTION

Post-transcriptional modification of RNA has in recent years emerged to be an important regulatory mode of gene expression. Over 150 distinct types of modifications have been found to decorate cellular RNAs, of which ribosomal RNA (rRNA) and transfer RNA (tRNA) are the most heavily modified (1). All four RNA bases and the ribose sugar can be targeted for modification, expanding the chemical properties of RNA nucleotides and hence, directly influencing RNA structure (2), stability (3), splicing (4), cellular localization (5) and interactions with RNA-binding proteins (2, 6). The recent findings of the dynamicity and reversible nature of RNA modifications have highlighted their involvement in crucial biological processes, such as development (7), cell fate determination (8), pathogen infection (9–11) and stress response (12, 13).

C5 methylation of cytosines (5-methylcytosine or m^5^C) is a common modification found on a wide variety of RNA molecules that ranges from ribosomal RNAs (rRNAs), transfer RNAs (tRNAs), messenger RNAs (mRNAs), vault RNAs (vtRNAs) to divers non-coding RNAs (5,14–17). In eukaryotes, m^5^C modification on RNA is catalyzed by the members of the NOL1/NOP2/SUN domain family of S-adenosylmethionine (SAM)-dependent methyltransferases (NSUN1 to 7), as well as the DNA methyltransferase homolog DNMT2. NOP2/NSUN1, NSUN2, and NSUN5 are conserved in all eukaryotes, while the other family members (NSUN3/4/6/7) are only found in higher eukaryotes (3). The biological impact and extent of RNA cytosine methylation are not entirely understood. However, the functional characterizations of NSUN proteins and the identification of their RNA substrates have helped uncover several essential roles played in the regulation of nuclear and mitochondrial gene expression (14,18–20). Mutations and changes in expression levels of genes coding for several NSUN proteins have been linked with diverse human diseases ranging from neurological disorders (21) to cancer (22–24), highlighting the importance of further characterizing this family of RNA methyltransferases.

NOP2/NSUN1 was initially identified in humans as p120; a proliferating-cell nucleolar antigen found overexpressed in a variety of rapidly dividing cells and malignant cancers (22,23,25–35). NOP2/NSUN1 also appears to harbor oncogenic function as its overexpression is sufficient to increase proliferation and confer tumorigenic potential *in vivo* (36). Studies in budding yeast have shown that NOP2/NSUN1 is an essential protein required for pre-rRNA processing and 60S ribosome subunit synthesis. In *S. cerevisiae*, the NOP2/NSUN1 homolog has been shown to catalyze m^5^C deposition on the 25S rRNA at positions 2870, which lie near the peptidyl-transferase center of the ribosome (37). m^5^C has been proposed to stabilize rRNA folding for the formation of this functional domain, as it promotes base stacking and increases the stability of hydrogen bonds with guanine (38). An equivalent cytosine at position 4447 in human 28S rRNA is known to be methylated (39, 40) and NOP2/NSUN1 is predicted to catalyze this modification based on homology and functional complementation experiments in yeast (41). Recent studies in *C. elegans* have suggested, but not directly demonstrated, that NOP2/NSUN1 methylates the 26S rRNA at position C2982, which is predicted to be the equivalent of human C4447 (42). Although NOP2/NSUN1-dependent deposition of m^5^C into rRNAs has been assessed in budding yeast and worms, little is known about the roles and substrates of NOP2/NSUN1 in human cells.

Here, we utilized miCLIP-seq (methylation individual-nucleotide-resolution crosslinking and immunoprecipitation-sequencing) to identify the methylated targets of NOP2/NSUN1 in human cells. We find that rRNA is the major methylation-specific target of NOP2/NSUN1 and we demonstrate that it catalyzes the deposition of m^5^C at residue 4447 of the 28S rRNA as predicted. Interestingly, we also find that in addition to site-specific methylation of the 28S rRNA, NOP2/NSUN1 also regulates pre-rRNA processing through non-catalytic complex formation with box C/D snoRNAs. Our findings indicate that NOP2/NSUN1 is required to recruit U3 and U8 snoRNAs to the pre-90S ribosomal particle and that it facilitates their stable assembly into snoRNP complexes. Remarkably, we find that the m^5^C catalytic activity of NOP2/NSUN1 is not required for efficient rRNA processing, stable assembly of box C/D snoRNP complexes, and cell proliferation. Our study identifies for the first time the spectrum of RNA bound by NOP2/NSUN1 and reveals additional functions in ribosome biogenesis beyond catalyzing rRNA substrate methylation.

## MATERIAL AND METHODS

### Reagents

Reagents and kits used in this study were: Amersham Hybond N+ nylon membrane (RPN303B, GE healthcare), BCA kit (23225, Pierce), Biotinylated cytidine (bis)phosphate (NU-1706-Bio, Jena Bioscience), Immun-Star AP substrate (1705018, Bio-Rad), iTaq universal SYBR green SuperMix (1725125, Bio-Rad), Lipofectamine RNAi Max (13778075, Invitrogen), Monarch RNA cleanup kit (T2030L, NEB), Murine RNase inhibitor (M0314L, NEB), Nitrocellulose membrane (10600008, GE Healthcare), Protease inhibitors cocktail (78437, Thermo Scientific), Protein G Dynabeads (10003D, Invitrogen), Protein G agarose beads (101242, Invitrogen), Q5 2X master mix (M0492S, NEB), Q5 Site-Directed Mutagenesis Kit (E0554, NEB), RevertAid reverse transcriptase (EP0441, Thermo Scientific), Streptavidin-alkaline phosphatase conjugate (434322, Invitrogen), Streptavidin-IR800 (926-32230, LI-COR), SuperScript III (18080044, Invitrogen), Taq polymerase (M0273S, NEB), Random hexamer primers (SO142, Thermo Scientific), Thiazolyl blue tetrazolium bromide (MTT, T3450, Biosynth), TriZol reagent (15596018, Invitrogen), TurboDNase (AM2239, Invitrogen), Puromycin (P8833, Sigma), Protease K (P8107S, NEB). Antibodies used were: NOP2 (A302-018A, Bethyl), FLAG (F1804, Sigma), FBL (A303-891A, Bethyl), NOP56 (PA5-78329, Invitrogen), 15.5K (PA5-22010, Invitrogen), La protein (A303-901A, Bethyl), RUVBL1 (A304-716A, Bethyl), NUFIP1 (A303-148A, Bethyl), p53 (SC-126, Santa Cruz Biotechnology), p21 (556430, BD Biosciences), GAPDH (47724, Santa Cruz Biotechnology), Puromycin (PMY-2A4-s, DSHB), Normal rabbit IgG (NI01, Millipore).

### Biological resources

HCT116 wildtype (WT), HCT116 p53-/-, and HEK293T were purchased from ATCC. Lentiviral plasmid empty backbone pLKO-Tet-On was obtained from Addgene (21915). pLenti-CMV-GFP-Puro plasmid was obtained from Addgene (17448). Plasmid p3XFLAG -CMV-7.1 was obtained from Sigma (E7533).

### Data Availability/Sequence Data Resources

The datasets generated and/or analyzed during the current study are publicly available at the GEO genomics data repository (accession GSE188735).

### Cell culture and proliferation assay

HCT116 wildtype (WT), HCT116 p53-/-, and HEK293T cells were maintained in high glucose Dulbecco’s modified Eagle’s medium (DMEM) with 5% fetal bovine serum (FBS), 100 U/ml penicillin, and 100 µg/ml streptomycin under 5% CO_2_ atmosphere at 37 °C. Cells were replated into 48-well plates at 10 000 cells per well 48 h after siRNA transfection. From 72 h to 168 h after siRNA transfection, cell counts per well were determined using an MTT assay. MTT reagent was added to the culture at 0.5 mg/ml and incubated for 4 h. Media was carefully removed and MTT formazan crystals were dissolved by DMSO. MTT formazan concentration, which correlates with cell number, was read at 570 nm.

### RNA interference

Synthetic short interfering RNA (siRNA) oligonucleotides (Sigma) were delivered into cells using Lipofectamine RNAi Max with the reverse transfection protocol according to the manufacturer’s instructions. Briefly, 20 nM of siRNA oligonucleotide was incubated with 6 μl of Lipofectamine RNAi Max in a 6-well plate format. Cell suspension (150 000 cells/well) was then added to the well containing the transfection mixture. The following siRNA sequences were used: Control (non-targeting scramble): 5’-GAUCAUACGUGCGAUCAGATT-3’, siNOP2#1: 5’-CACCUGUUCUAUCACAGUATT-3’, siNOP2#2: 5’-GCAACGAUCACCUAAAUUATT-3’.

### Lentivirus construction and virus production

NOP2 shRNA sequences (oligo sequence available in Supplementary Table 1) were cloned into pLKO-Tet-On as described by Wiederschain *et al* (43). To overexpress NOP2/NSUN1 for rescue assays, the GFP gene in pLenti-CMV-GFP-Puro was replaced with siRNA resistant NOP2/NSUN1 cDNA or corresponding catalytically inactive version (C513A). Synonymous mutations for siRNA resistant and C513A mutation were created using the NEB Q5 Site-Directed Mutagenesis Kit and the primers utilized are listed in Supplementary Table 1. Lentivirus was produced by co-transfecting HEK-293T cells with lentiviral vectors (pLKO-Tet-On-shNop2 or pLenti-CMV-NOP2-Puro), pMDL, pVSVG, pREV in a ratio of 3:1:1:1 (44). Media was renewed 24h after transfection and virus-containing media was collected 48h and 72h after transfection.

### Exogenous expression of wildtype and C459A mutant NOP2

Wildtype NOP2/NSUN1 gene was cloned into p3XFLAG -CMV-7.1 plasmid. The C459A mutation was created using the NEB Q5 Site-Directed Mutagenesis Kit and the primers listed in Supplementary Table 1. Sequences of NOP2/NSUN1 WT and C459A constructs were confirmed by Sanger sequencing (Genewiz).

### RNA extraction and Northern blot analysis

Total RNA was extracted using TriZol reagent following the manufacturer’s instructions. 2.5 µg of total RNA was separated on 0.8% formaldehyde denaturing agarose gel or 10% 8M urea-PAGE gel and transferred to Amersham Hybond N+ nylon membrane. RNA was crosslinked to membrane by 120 mJ/cm^2^ UV exposure. Membrane was pre-hybridized in hybridization buffer (0.6 M NaCl, 60 mM sodium citrate pH 7, 0.5% SDS, 0.1% BSA, 0.1% polyvinyl pyrrolidone, 0.1% ficoll 400) at 50 °C for 1h and hybridized with 20nM biotin-labeled probe in fresh hybridization buffer overnight. Probe was detected using streptavidin-alkaline phosphatase conjugate and Immun-Star AP substrate. Probes for Northern blot are listed in Supplementary Table 1. ImageJ was used for densitometry quantification analysis of each blot.

### RNA immunoprecipitation (RNA-IP) and RT-qPCR

To assess NOP2/NSUN1, 15.5K, or Nop56 association to snoRNAs by RNA-IP, cells were lysed in lysis buffer (1% NP-40, 50 mM Tris-HCl pH 8, 150 mM NaCl, 5 mM EDTA, 100 U/ml murine RNase inhibitor, 1X protease inhibitors cocktail) on ice for 15 min and triturated 10 times through a 27-gauge needle. The lysates were cleared by centrifugation at 15000 g for 10 min at 4 °C and the supernatant was incubated with 10 μl of protein G Dynabeads pre-bound with 1 μg of antibody or normal IgG. After 2 h incubation, beads were washed 5 times with cold high salt wash buffer (1% NP-40, 50 mM Tris-HCl pH 8, 500 mM NaCl, 5 mM EDTA) and 2 times with no salt wash buffer (20 mM Tris-HCl pH 7.4, 0.2% Tween-20). Beads were treated with DNase followed by proteinase K. Immuno-precipitated RNA was extracted using TriZol reagent. RNA samples were reverse transcribed using RevertAid reverse transcriptase and random hexamer. qPCR was performed with 500 nM specific primer mix and iTaq universal SYBR green SuperMix on a C1000 Touch CFX-384 real-time qPCR system (Bio-Rad). Primer efficiency test was performed using serial dilution of cDNA made from 1 μg total RNA. Efficiency was calculated using the formula: efficiency = 10 ^ (−1/Slope of the standard curve). Primers for qPCR are listed in Supplementary Table 1.

### Immunoprecipitation and Immunoblotting

For co-immunoprecipitation analysis, cells were lysed in ELB buffer (0.5% NP-40, 50 mM HEPES pH 7.2, 250 mM NaCl, 2 mM EDTA, 1X protease inhibitors cocktail) on ice for 15 min. The lysates were cleared by centrifugation at 15000 g for 10 min at 4 °C and the supernatant was incubated with 20 μl of protein G agarose beads pre-bound with 1 μg of antibody or normal IgG. After 2 h incubation, beads were washed 4 times with ELB buffer and re-suspended in 15 μl 2X protein loading dye (4% SDS, 0.25 M Tris-HCl pH 6.8, 20% glycerol, 3% beta-mercaptoethanol, 0.04% bromophenol blue). For immunoprecipitation of RUVBL1, NUFIP1, and La protein, the following protocol was used. Cells were lysed in lysis buffer (1% NP-40, 50 mM Tris-HCl pH 7.5, 150 mM NaCl, 2 mM MgCl_2_, 1X protease inhibitors cocktail) on ice for 15 min. The lysates were sonicated in pulses of 1 second, 12 times on a Misonix XL-2000 sonicator at power intensity 3 and cleared by centrifugation at 15000 g for 10 min at 4 °C. The supernatant was incubated with 20 μl of protein G agarose beads pre-bound with 2 μg of antibody or normal IgG. After 2 h incubation, beads were washed 4 times with lysis buffer and re-suspended in 15 μl 2X protein loading dye. For total protein extraction, cells were washed with PBS, harvested by scraping, and lysed in RIPA buffer (25mM Tris-HCl pH 7.6, 150mM NaCl, 1% NP-40, 1% Triton X-100, 1% sodium deoxycholate, and 0.1% SDS) plus protease inhibitors cocktail for 15 min on ice. Lysates were cleared by centrifugation at 22,000 x g for 10 min at 4°C. Protein concentrations were evaluated with the BCA kit. Proteins were separated on SDS-poly-acrylamide gel electrophoresis (SDS-PAGE) and transferred to a nitrocellulose membrane. ImageJ was used for densitometry quantification analysis.

### Nuclear and nucleolar extraction for immunoprecipitation

Nuclei were isolated from HCT116 cells as described by Pestov *et al* (45). Nuclear and nucleolar extracts used for co-immunoprecipitation were prepared as described by Watkins *et al* (46) with minor modifications. Briefly, isolated nuclei were re-suspended in DM buffer (20 mM HEPES-NaOH pH 7.9, 150 mM NaCl, 3 mM MgCl_2_, 0.2 mM EDTA, 0.5 DTT, 10% Glycerol, 1X protease inhibitors cocktail) and sonicated in pulses of 1 second, 12 times on a Misonix XL-2000 sonicator at power intensity 3. The lysates were cleared by centrifugation at 16000 g for 30 min at 4 °C. NP-40 and EDTA were added to the supernatants to final concentrations of 1% and 5 mM respectively, and the resulting lysate was used as nuclear extract. To prepare nucleolar extracts, the pellets were re-suspended in buffer D (DM buffer without MgCl_2_) and sonicated in pulses of 1 second, 20 times on a Misonix XL-2000 sonicator at power intensity 3. The lysates were cleared by centrifugation at 16000 g for 30 min at 4 °C. NP-40 and EDTA were added to the supernatants to final concentrations of 1% and 5 mM respectively, and the resulting lysate was used as nucleolar extract. 1X protein loading dye (2% SDS, 0.12 M Tris-HCl pH 6.8, 10% glycerol, 1.5% beta-mercaptoethanol, 0.02% bromophenol blue) was added to the pellet and sonicated in pulses of 1 second, 10 times on a Misonix XL-2000 sonicator at power intensity 3. The mixture was cleared by centrifugation at 25000 g for 10 min at 4 °C. The supernatant containing insoluble nucleolar proteins was used for Western blot. IP-WB and RNA-IP experiments using nuclear and nucleolar extraction were done as described in “Immunoprecipitation and Immunoblotting” and “RNA immunoprecipitation (RNA-IP) and RT-qPCR” section respectively.

### Immunofluorescence staining

To localize exogenous NOP2/NSUN1 by immunofluorescence, HCT116 cells expressing FLAG-tagged WT or C459A mutant NSUN1/NOP2 were grown on cover glass, fixed with 4% formaldehyde, immunostained with anti-FLAG and anti-FBL antibodies, and visualized by fluorescence microscopy as described in our previous paper (47). Similarly, localization of endogenous NOP2/NSUN1 was performed with an anti-NOP2/NSUN1 antibody from Bethyl (A302-018A).

### Polysome preparation

HCT116 cells were infected with doxycycline-inducible NOP2/NSUN1 shRNA-expressing lentivirus. After puromycin selection, cells were treated with 200 ng/ ml doxycycline (Dox) for 4 days before harvesting to induce shRNA expression. Non-Dox-treated cells were used as control. Polysome profile was done as described previously by Simsek *et al.,* (48). In brief, cells were treated with 100 μg/ml cycloheximide for 5 min and lysed in polysome buffer (25 mM Tris-HCl pH 7.5, 150 mM NaCl, 15 mM MgCl_2_, 8% glycerol, 1% Triton X-100, 0.5% sodium deoxycholate, 1 mM DTT, 100 ug/ml cycloheximide, 100 U/ml murine RNase inhibitor (NEB), 25 U/ml TurboDNase, 1X protease inhibitor cocktail). The lysate was cleared by a serial of centrifugation, loaded on 10%-50% sucrose gradient, and ultra-centrifuged at 40 000 RPM for 2.5h at 4°C with a Beckman Ti-41 swing rotor. Fractions were collected using a piston gradient fractionator (BioComp).

### Sucrose gradient density centrifugation for co-sedimentation assays

To prepare core ribosomal particles and nuclear extracts, nuclei from cells were extracted first as described by Pestov *et al.,* (45). Nuclear extracts were prepared by sonication of nuclei in sonication buffer (25 mM Tris-HCl pH 7.5, 100 mM KCl, 2 mM EDTA, 1 mM NaF, 0.05% NP-40) and then centrifugation at 15000 g for 15 min at 4°C. Core preribosomal particles were extracted from nuclei as described in Lapik *et al*., (49). Lysates of nuclear extracts and core preribosomal preparations were loaded on 10%-30% sucrose gradient, and ultra-centrifuged at 36000 RPM for 3h at 4°C with a Beckman Ti-41 swing rotor. Fractions were collected using a piston gradient fractionator (BioComp). Proteins from each fraction were extracted as described by Pestov *et al.,* (45) for Western blot analysis. To isolate RNA, fractions were digested with 3.2 U/ml protease K, 1% SDS, and 5 mM EDTA at 42°C for 1 h, and extracted with phenol-chloroform.

### Puromycin labeling assay

Seventy-two hours after siRNA transfection, HCT116 WT cells were replated into 6-well plate at 8 x 10^5^ cells/well and incubated for another 12 h. Cells were treated with 5 μg/ml puromycin for 0, 15, and 30 min and immediately harvested for Western blot analysis using an anti-puromycin antibody.

### Biotin-labelling of immunoprecipitated RNA

pCp-Biotin labeling of immunoprecipitated RNA was performed according to (50). Briefly, four million HEK293T cells expressing empty vector, FLAG-NOP2-WT or FLAG-NOP2-C459A were lysed in CLIP buffer (50 mM Tris-HCl pH 7.4, 100 mM NaCl, 1% NP-40, 0.1% SDS, 0.5% sodium deoxycholate, 1X Protease Inhibitor Cocktail) without UV crosslink on ice for 15 min followed by sonication and digestion with RNase T1 and Turbo DNAse at 37°C for 5 min. Digested lysates were cleared by centrifugation and immunoprecipitated using 25 μl of protein G Dynabeads pre-bound with 2 μg of anti-FLAG antibody at 4°C overnight. Samples were washed twice in high salt wash buffer (50 mM Tris-HCl pH 7.4, 1 M NaCl, 1% NP-40, 0.1% SDS, 0.5% sodium deoxycholate, 1 mM EDTA), thrice in wash buffer (20 mM Tris-HCl pH 7.4, 0.2% Tween-20) and RNA was dephosphorylated on beads by Fast-AP and T4 PNK and ligated with biotinylated cytidine (bis)phosphate (pCp-Biotin) using T4 RNA ligase. Samples were loaded on 8% SDS-PAGE gel for separation and transferred to a nitrocellulose membrane. Streptavidin-IR800 was used for detecting biotin-labeled RNA and the membrane was scanned using LI-COR Odyssey DLx Imager.

### miCLIP-sequencing

The eCLIP-sequencing protocol described by Van Nostrand *et al* (51) was applied with minor modifications. Twenty million cells were lysed in CLIP buffer (50 mM Tris-HCl pH 7.4, 100 mM NaCl, 1% NP-40, 0.1% SDS, 0.5% sodium deoxycholate, 1X Protease Inhibitor Cocktail) without UV crosslink on ice for 15 min followed by sonication and digestion with RNase T1 and Turbo DNAse at 37°C for 5 min. Digested lysates were cleared by centrifugation and immunoprecipitated using 100 μl of protein G Dynabeads pre-bound with 10 μg of anti-FLAG antibody at 4°C overnight. Samples were washed twice in high salt wash buffer (50 mM Tris-HCl pH 7.4, 1 M NaCl, 1% NP-40, 0.1% SDS, 0.5% sodium deoxycholate, 1 mM EDTA), once in wash buffer (20 mM Tris-HCl pH 7.4, 0.2% Tween-20) and RNA was dephosphorylated, treated with T4 PNK and ligated to 3’ RNA adapter (/5Phos/rArGrArUrCrGrGrArArGrArGrCrArCrArCrGrUrC/3SpC3/) on beads. Samples (including SMinputs) were loaded on 8% SDS-PAGE gel for separation, transferred to a nitrocellulose membrane and RNA was isolated from membrane by cutting a region 75 kDa above NOP2/NSUN1. CLIP and SMinput libraries were prepared as paired-end high-throughput libraries as described in (51) and sent for sequencing on Illumina Hi-Seq 4000 platform (Novogene). Raw sequencing data were adaptor trimmed using CutAdapt (52). A unique molecular identifier (UMI) was appended to the read name using fastp (53). Reads less than 18 nt were discarded. Trimmed reads were aligned to human genome (hg38) and human repetitive elements using family-aware repetitive elements mapping pipeline described by Van Nostrand *et al* (*50*). miCLIP peak calling was performed on mapped reads from the above repetitive elements mapping pipeline using Peakachu (https://github.com/tbischler/PEAKachu) with parameter --paired_end -- max_insert_size 220. To determine crosslink sites on rRNA, trimmed reads were aligned to 47S rRNA (NR_046235) using Bowtie2 (54) with parameter --no-mixed-a, and deduplicated using UMI-tools (55) with parameter --read-length. Mapping statistics, sequencing depth and read start were obtained using Samtools package (56) and reverse transcription (RT) stops were assigned at the start (+1) sites of Read1 sequence.

### Bisulfite-sequencing

Two micrograms of nuclear RNA were incubated with 2.3 M Na_2_S_2_O_5_ and 0.57 mM hydroquinone at 70°C for 5 min followed by 54°C for 75 min. Converted RNA was extracted using NEB Monarch RNA extraction kit and desulfonated with 1M Tris-HCl pH9 at 37°C for 1h. RNA was then precipitated in ice-cold ethanol. cDNA was made with superscript III reverse transcriptase and random 9-mer. PCR amplification of the 28S rRNA fragment from position 4178 to 4534 or the full length vtRNA1.2 was performed using NEB Taq polymerase and target-specific primers with Illumina adaptor overhangs. The PCR product was purified by gel extraction and sequenced on Mi-Seq platform with 2×250 paired-end mode (Genewiz EZ Amplicon sequencing, 130 000 reads/sample were aligned to 28S rRNA on average). The primers used for bisulfite sequencing are listed in Supplementary Table 1. Sequencing reads were aligned to the converted reference sequence using Bowtie2 with default settings. Mapping information was retrieved using Samtools package as described previously by Heng (56).

### Statistics

Data were shown as mean ± standard deviation. Each dot on dot-bar plot represents an independent biological replicate. The difference between two groups was determined by two-tailed Student’s *t*-test.

## RESULTS

### NOP2/NSUN1 binds to rRNA, vtRNA1.2 and snoRNAs

We utilized a miCLIP-sequencing approach to identify NOP2/NSUN1 RNA substrates and provide nucleotide resolution of the C5-methylated cytosines (16). A characteristic of NSUN methyltransferases is the presence of two catalytic cysteines in the active site. Preceding methylation, a covalent intermediate is first formed between motif VI cysteine (C513 on NOP2/NSUN1) and the RNA cytosine pyrimidine ring. After catalysis, the release of the methylated RNA depends on the second conserved cysteine located in motif IV (C459 on NOP2/NSUN1) (Figure 1A) and mutation of this cysteine has been shown to result in a stable covalent protein-RNA intermediate (3,57,58). miCLIP takes advantage of this particularity and allows NSUN proteins with a mutated cysteine (motif IV) to be immunoprecipitated with a covalently trapped RNA substrate without the need for UV crosslinking (Figure 1B) (16). Mutation of NOP2/NSUN1 conserved motif IV cysteine (C459A) resulted in the stabilization of covalently linked RNA intermediates as evidenced by a shift in molecular weight of immunoprecipitated FLAG-tagged C459A mutant expressed in HEK 293T cells compared with WT (Figure 1C). Ligation-mediated labeling of immunoprecipitated FLAG-NOP2/NSUN1 WT and C459A expressed in HEK 293T cells with pCp-Biotin (Cytidine-5’-phosphate-3’-(6-aminohexyl) phosphate labeled with Biotin), followed by detection with fluorescence imaging of streptavidin-conjugated-IR800 confirmed the presence of trapped protein-RNA complexes (Figure 1D, high exposure panel). Although not to the same extent as for the mutant, immunoprecipitated FLAG-WT also showed RNA labeling above control level, suggesting that NOP2/NSUN1 may be tightly bound to a subpopulation of RNAs (Figure 1D, densitometry analysis of high exposure panel). Immuno-fluorescence microscopy confirmed that both FLAG-NOP2/NSUN1 WT and C459A mutant localized to the nucleolus, demonstrating their suitability for miCLIP assays (Supplementary Figure S1A).

**Figure 1.**
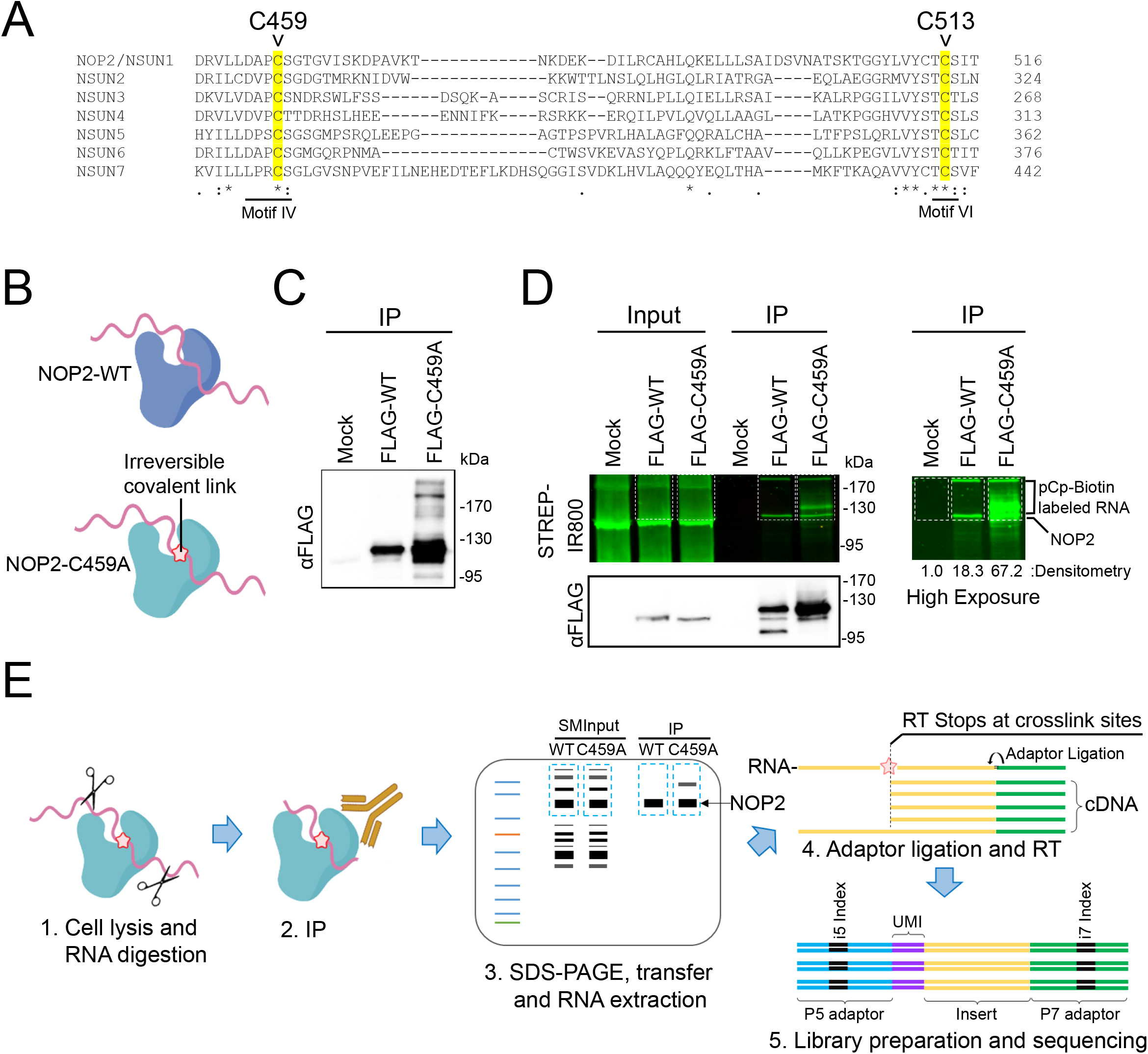
Human NOP2/NSUN1 miCLIP design. **(A)** Alignment of the protein sequences of human NSUN family members showing conserved cysteine residues (highlighted) required for catalytic activity (C459 and C513). The cysteine mutated for the miCLIP assay is C459 in motif IV **(B)** Cartoon depicting that only the NOP2/NUSN1 C459A mutant forms irreversible covalent crosslinks with RNA substrates. **(C)** FLAG-tagged NOP2/NSUN1 Wildtype (WT) and C459A mutant were expressed in HEK293T cells and immunoprecipitated with a FLAG antibody. Immuno-precipitated proteins were detected by Western Blotting using a FLAG antibody. **(D)** FLAG-tagged NOP2/NSUN1 WT and C459A mutant expressed in HEK293T cells were immunoprecipitated with a FLAG antibody. Co-precipitated RNA was 3’-end ligated with pCp-biotin for visualization. After SDS-PAGE separation and membrane transfer, the pCp-biotin-labeled RNA was detected with Streptavidin-IR800. Immunoprecipitated NOP2/NSUN1 WT and C459A mutant were detected by immunoblotting with a FLAG antibody. The right panel shows a high exposure of the pCp-biotin-labeled RNA with relative densitometry analysis. **(E)** Schematic overview of the NOP2/NSUN1 miCLIP-sequencing. HEK293T cells expressing NOP2/NSUN1 WT and C459A mutant were lysed, digested with RNase, immunoprecipitated with FLAG antibody, separated on SDS-PAGE and transferred to membrane. NOP2/NSUN1 WT and C459A mutant-associated RNAs and their respective size-matched inputs (SMInputs) were extracted from the membrane and processed for libraries and Illumina sequencing.

Following limited RNase digestion and immunoprecipitation with a FLAG antibody, co-precipitated RNA from Mock, WT and C459A mutant complexes from HEK 293T cells (as well as their respective paired size-matched input (SMInput)) were purified by separation on SDS-PAGE and then transferred onto membrane. Protein-RNA complexes from a 75 kDa region above NOP2/NSUN1 were excised from membrane (Figure 1E). Recovered RNA was further prepared into paired-end libraries for high-throughput sequencing on Illumina Hi-Seq 4000 platform. Only samples from biological duplicates of WT, C459A mutant and respective paired SMInput were processed for sequencing as the Mock immunoprecipitated samples did not recover enough RNA to be amplified for library preparation. Reads were processed using a modified eCLIP-seq pipeline (50). The numbers of reads identified as binding sites in miCLIP were normalized to SMInput and SMInput-normalized significance and fold enrichment was calculated (Supplementary Table 2).

The majority of miCLIP reads for the C459A mutant (>97%) corresponded to rRNA (Figure 2A), which also showed peaks of significant enrichments when normalized to SMInput (Figure 2B). The remaining reads mostly mapped to different types of non-coding RNAs (ncRNA) where significantly enriched peaks corresponded to vtRNA1.2 and several snoRNAs (1.1%), with U3 (SNORD3) and U8 (SNORD118) being some of the most represented (Figure 2B). NOP2/NSUN1 C459A mutant bound primarily to box C/D snoRNAs (SNORD), including U3 and U8. Only a few H/ACA box snoRNAs were found, indicating a specificity of interaction for NOP2/NSUN1 with box C/D snoRNAs (Figure 2C, Supplementary Table 2). Surprisingly, most of the reads from WT NOP2/NSUN1 also mapped to rRNA (>98%), followed by C/D box snoRNAs and various ncRNAs (Figure 2A-C, Supplementary Table 2). vtRNA1.2 was only recovered cross-linked to the C459A NOP2/NSUN1 mutant and did not show any significant enrichment in the WT IP, suggesting vtRNA1.2 could be a catalytic substrate (Figure 2B, Supplementary Table 2). To confirm the interaction with box C/D snoRNAs, we independently immunoprecipitated FLAG-NOP2/NSUN1 WT and C459A mutant in miCLIP conditions but without membrane transfer followed by qRT-PCR analysis. Both WT and mutant showed significant binding enrichment to box C/D snoRNAs but not to U6 snRNA and the H/ACA box snoRNA SNORA7A, indicating specificity of interaction (Supplementary Figure S1B). Together, these findings confirm that NOP2/NSUN1 likely forms strong interactions with box C/D snoRNAs that resist the stringent wash conditions utilized during the CLIP purification.

**Figure 2.**
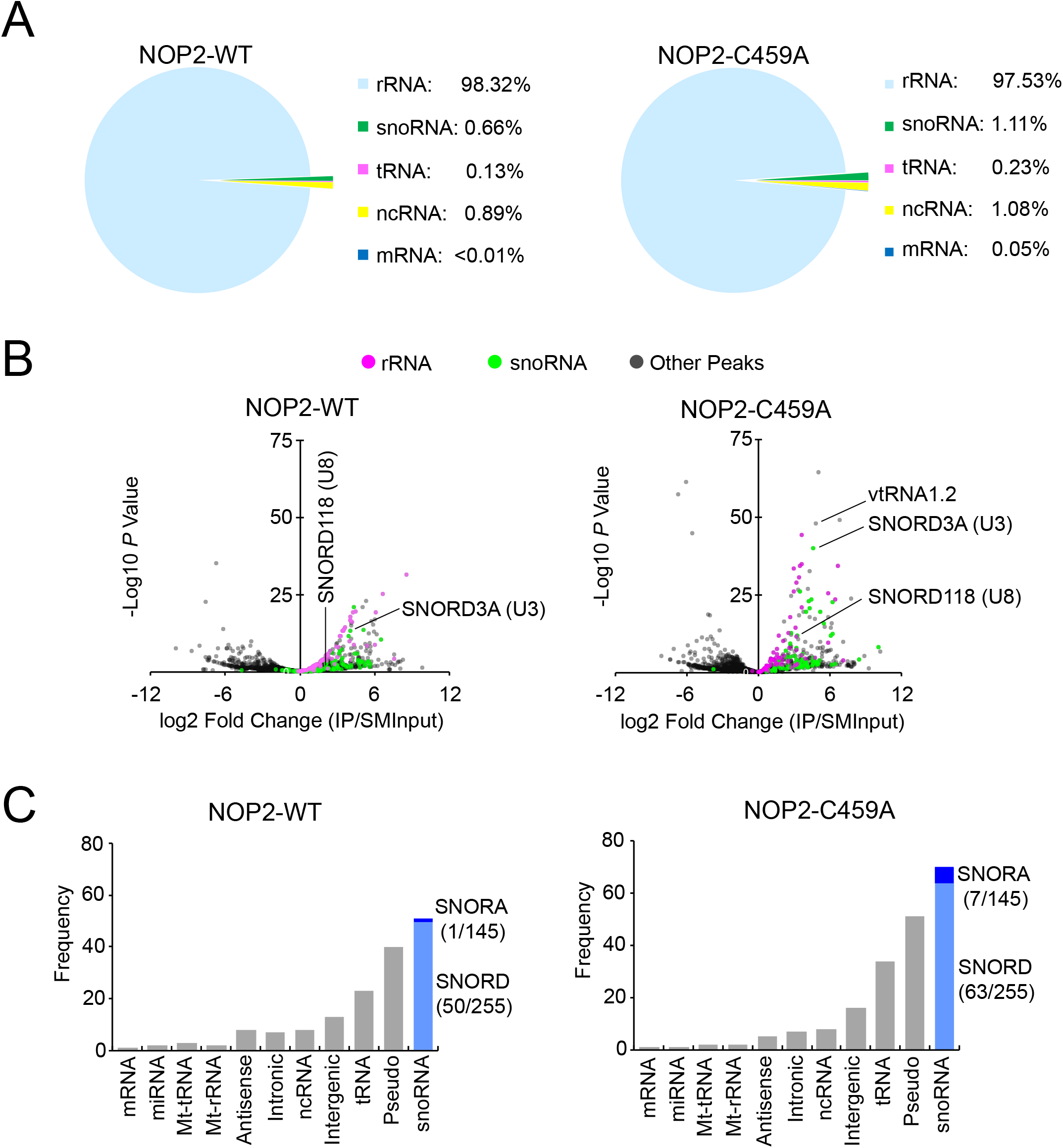
Human NOP2/NSUN1 binds to rRNA and snoRNAs. **(A-C)** HEK293T cells expressing FLAG-tagged NOP2/NSUN1 WT (non-covalently bound to substrate) or C459A mutant (covalently bound to substrate) were subject to CLIP-sequencing analysis (two biological replicates for each sample). CLIP-sequencing reads were aligned to human genome (hg38) and repetitive elements families. CLIP peaks were called by Peakachu. **(A)** Percentage of miCLIP mapped peak regions in each indicated RNA category with cut-offs of adjusted significance *P* values < 0.05 and fold change over size-matched input (SMInput) >= 2. **(B)** Volcano plots of all peaks detected by Peakachu. Each dot represents a peak either enriched in IP (Log2 fold change > 0) or SMInput samples (Log2 fold change < 0). **(C)** Number of CLIP peaks with cut-offs of adjusted significant *P* values < 0.05 and fold change over SMInput >= 2 in each indicated RNA category. C/D box and H/ACA box snoRNA are shown in light and dark blue, respectively. Number of snoRNAs found in CLIP peaks / total number of snoRNAs in the indicated subfamily are shown in parentheses.

### NOP2/NSUN1 methylates the 28S rRNA at position C4447

Human rRNA is transcribed by RNA Pol I as a long precursor transcript (47S) that undergoes a complex series of cleavage and modification steps to generate the mature 18S, 28S, and 5.8S rRNAs (59, 60). 5S rRNA is separately transcribed by RNA Pol III (61). Alignment of the CLIP reads to the 47S precursor rRNA revealed that both NOP2/NSUN1 WT and C459A mutant have a similar and specific binding pattern to the 5’ external transcribed spacer (5’ETS) region encompassing the 01/A’, A0 and 1 endonucleolytic processing cleavage sites (Figure 3A). Interestingly, only the C459A mutant showed a distinct enrichment peak on the 28S rRNA that mapped exactly at C4447, which corresponds to the predicted NOP2/NSUN1 methylation site based on homology to the yeast counterpart (Figure 3A). The NOP2/NSUN1 C459A mutation causes irreversible covalent crosslinks between the protein and target cytosine of the RNA substrate, causing the reverse transcription (RT) to terminate precisely at the polypeptide-cytosine 5 crosslink site (16). To identify NOP2/NSUN1 target cytosines, RT stops were mapped and m^5^C were assigned at the +1 site of sequencing reads. A clear enrichment of reads starting at C4447 on the 28S rRNA was observed only for the C459A mutant (Figure 3B), suggesting that this site is methylated by NOP2/NSUN1. Examination of all CLIP reads identified only one additional cross-link site at C27 of vtRNA1.2, which we found to be specific to the C459A mutant only (Figure 3C). Interestingly, C27 on vtRNA1.2 has previously been identified as an NSUN2 target site by miCLIP (16), suggesting that both NSUN2 and NOP2/NSUN1 could regulate the function of vtRNA1.2 through m^5^C modification. Our analysis did not identify potential methylations or cross-linked sites on snoRNAs (Supplementary Figure S2A-B). We found that WT NOP2/NSUN1 and the C459A mutant bound to similar regions of mature box C/D snoRNAs, including U3 and U8 (Supplementary Figure S2C-D), suggesting that the interaction with box C/D snoRNAs is likely independent of NOP2/NSUN1 methyltransferase activity.

**Figure 3.**
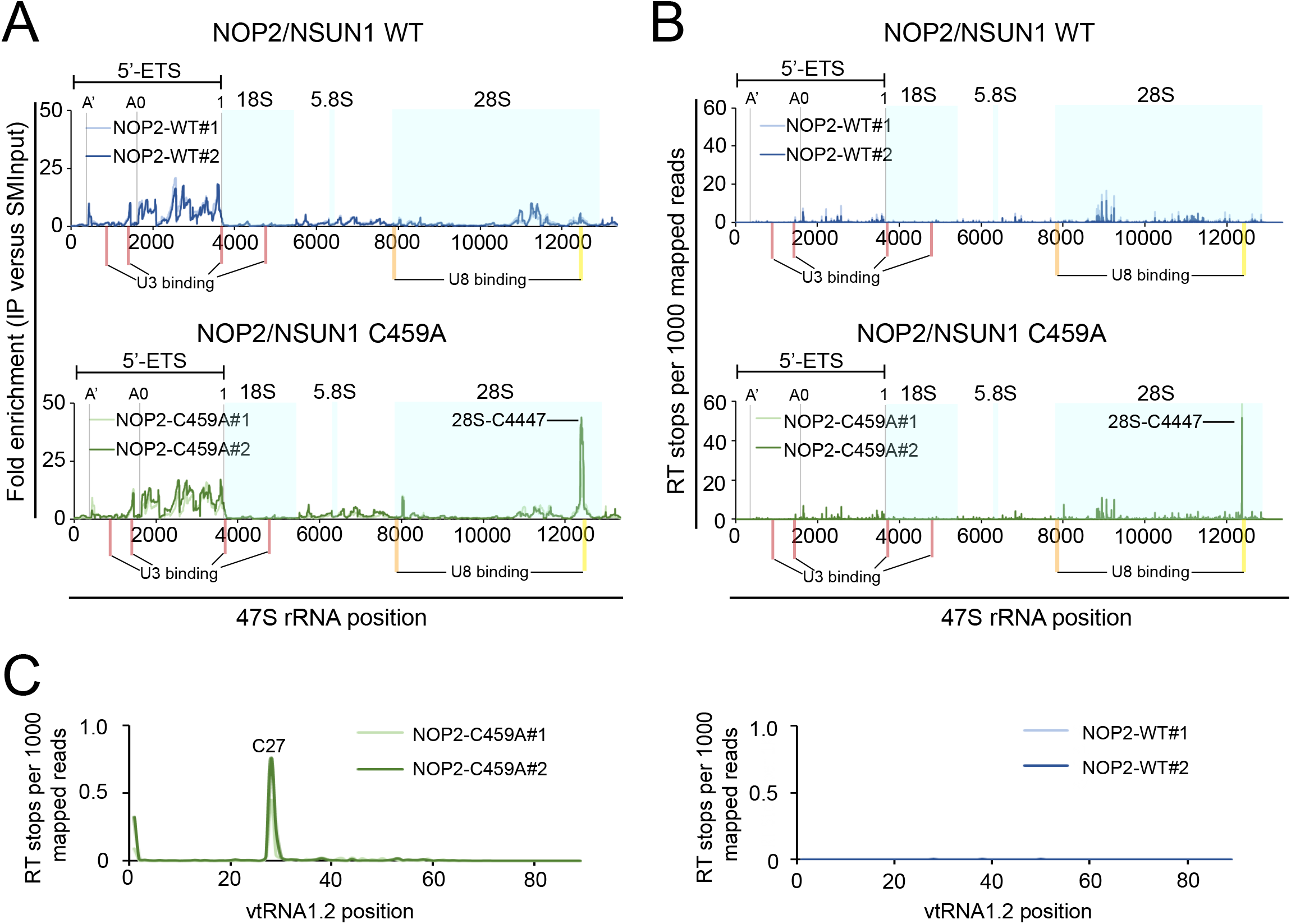
Human NOP2/NSUN1 binds to the rRNA 5’-ETS region and crosslinks on 28S rRNA at position C4447. **(A-B)** miCLIP-sequencing data were aligned to human 47S pre-rRNA (NR_046235) and the mapping information was retrieved using Samtools. The location of mature 18S, 5.8S, and 28S rRNAs are marked with light blue shadow boxes. A’, A0 and 1 cleavage sites on 5’-ETS are marked with gray lines. Red bars indicate the location of U3 snoRNA binding sites (101). The orange bar indicates the location of U8 snoRNA binding site based on sequence homology and accessibility measurements (103). The yellow bar indicates the location of the U8 snoRNA binding site found by RNA duplex mapping (106). **(A)** Plot of sequencing depth-normalized reads ratio of IP over size-matched input (SMInput) on the 47S rRNA. **(B)** Plot of reverse transcription (RT) stops on the 47S rRNA normalized by sequencing depth. RT stops were assigned at the start (+1) sites of the Read1 sequence from NOP2/NSUN1 WT and C459A IP samples. **(C)** Plot of reverse transcription (RT) stops on vtRNA1.2 normalized by sequencing depth. RT stops were assigned at the start (+1) sites of the Read1 sequence from NOP2/NSUN1 WT and C459A IP samples. **(B-C)** Plotted data is represented for each independent biological replicate for NOP2/NSUN1 WT (blue lines) and C459A mutant (green lines).

To confirm the NOP2/NSUN1-mediated methylation sites identified by miCLIP on 28S rRNA and vtRNA1.2, we performed RNA bisulfite sequencing in cells treated with control vs. NOP2/NSUN1 siRNAs (Figure 4A). In line with the miCLIP data, a bisulfite-resistant cytosine resulting from C5 methylation was detected at positions C4447 of the 28S rRNA while all surrounding cytosines were efficiently deaminated to uracil (overall conversion rate of 97%) (Figure 4B). Because we used nuclear RNA to perform the bisulfite conversion, a small fraction (16%) of C4447 was detected as unmethylated, likely representing newly synthesized precursor rRNA prior to modification (Figure 4B-C). RNAi depletion of NOP2/NSUN1 resulted in a significant loss in bisulfite resistant (non-deaminated) cytosine at this position as shown by an increase to 28% (siRNA #1) or 34% (siRNA #2) of converted cytosine at position 4447 in the NOP2/NSUN1 knockdowns compared with control (Figure 4B-C). We estimate to have about 15% of residual NOP2/NSUN1 protein remaining after RNAi, which could explain why we do not observe a complete loss of m^5^C4447 in the NOP2/NSUN1 RNAi-treated samples. These findings confirm that NOP2/NSUN1 is responsible for the deposition of m^5^C at residue 4447 of the 28S rRNA. Because this conclusion is based on NOP2/NSUN1 knockdown with about 15% residual protein, the possibility that NOP2 is not the only methyltransferase responsible for the deposition of m5C at 4447 cannot be excluded.

**Figure 4.**
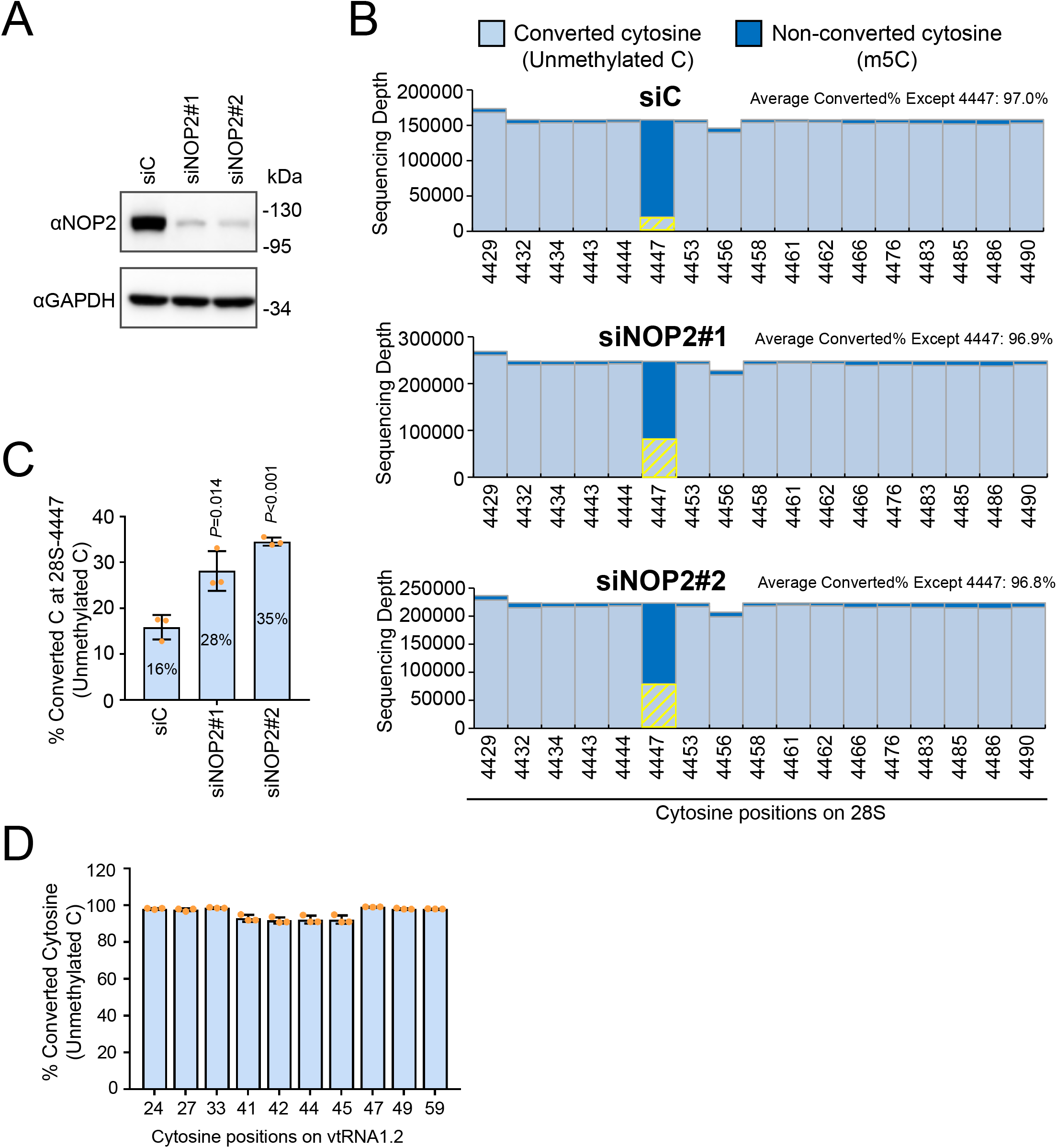
Human NOP2/NSUN1 methylates the 28S rRNA at position C4447. **(A)** HCT116 cells were transfected with non-targeting control (siC), NOP2/NSUN1 siRNA#1 or siRNA#2. 72h later, NOP2/NSUN1 depletion efficiency was determined by Western blot. **(B)** A fraction of cells from (A) were harvested for nuclear RNA extraction. Nuclear RNA was treated with bisulfite salt and the region surrounding C4447 was amplified by RT-PCR for sequencing on the Illumina Mi-Seq platform. Non-converted (protected by 5mC modification) and converted cytosines (lacking m5C modification) near C4447 were plotted. The converted cytosines at position 4447 are highlighted by yellow bars. **(C)** Percentage of converted cytosine (C) at position 4447 on 28S rRNA. **(D)** Percentage of converted cytosine on vtRNA1.2. Total RNA from HCT116 cells was treated with bisulfite salt and vtRNA1.2 was amplified by RT-PCR for sequencing on the Illumina Mi-Seq platform. The data are shown as the mean of 3 independent biological replicates ± standard deviation (SD). Statistical significance between NOP2/NSUN1 depleted samples and non-targeting siRNA control samples was calculated using a 2-tailed independent student *t*-test.

RNA bisulfite sequencing analysis did not find any m^5^C on vtRNA1.2 (Figure 4D), which is consistent with the study of Hussain *et al.* (16) that also identified C27 on vtRNA1.2 as an NSUN2 target by miCLIP but failed to detect any m^5^C by bisulfite sequencing. NSUN2-mediated m^5^C of vtRNAs has been shown to positively regulate their cleavage into specific small RNAs called vtRNA-derived small RNAs (svRNAs) (16, 62). It is possible that most of the vtRNA1.2 pool methylated at C27 is subjected to endonucleolytic processing and thus not amenable for detection by bisulfite sequencing. Further experiments will be needed to confirm whether the interaction between NOP2/NSUN1 and vtRNA1.2 is real and whether vtRNA1.2 C27 is a *bona fide* methylation target.

### NOP2/NSUN1 is required for efficient rRNA processing and 60S ribosome biogenesis

Our miCLIP analysis revealed non-crosslinked binding to the 5’ETS region of the pre-rRNA transcript and to box C/D snoRNAs for both WT and C459A mutant (Figure 2B and 3A-B), suggesting that NOP2/NSUN1 has additional functions beyond catalyzing target modification. Previous CLIP studies have found that RNA-binding proteins (RBPs) binding to U3 and U8 box C/D snoRNAs also had correlated binding enrichment to the 5’ETS of the rRNA precursor (50,63,64). U3 and U8 regulate folding and cleavage reactions leading to synthesis of the small and large subunit rRNAs, respectively (65–67). U3 depletion was shown to impair processing of the 5′ ETS and ITS1 regions. In contrast, depletion of U8 was found to primarily affect 3’ETS processing and decrease the levels of the 32S and 12S rRNA intermediates of the mature 28S and 5.8S rRNAs, respectively (65). To investigate whether NOP2/NSUN1 regulates U3 and U8 box C/D snoRNAs-dependent processing events, we first performed a detailed pre-rRNA processing analysis by Northern blotting using specific probes detecting most of the major pre-rRNA intermediates (Figure 5A). We find that depletion of NOP2/NSUN1 severely affected the maturation steps leading to the formation of the 5.8S and 28S rRNAs from the 60S subunit, as observed by a marked reduction in 32S and 12S intermediates (Supplementary Figure S3A-B and S4A). Processing of the 5’ETS and 3’ETS was also compromised as evidenced by the modest but consistent increase of the 47S, 45S and 41S transcripts and the accumulation of pre-rRNAs showing abnormal extension of retained 3′-ETS sequences (most evident for 41S-L and 36S-L). These observations indicate inefficient cleavage at sites A’, A0, 1 and 02 (Supplementary Figures S3A-B and S4A) and are similar to the processing defects observed upon U3 and U8 knockdowns (65). Together our findings suggest that, in addition to methylating the 28S rRNA at position C4447, NOP2/NSUN1 appears important for multiple steps of rRNA processing. These NOP2/NSUN1 functions are likely independent of its catalytic activity and linked to its binding to the 5’ETS regions and U3 and U8 snoRNAs (65).

**Figure 5.**
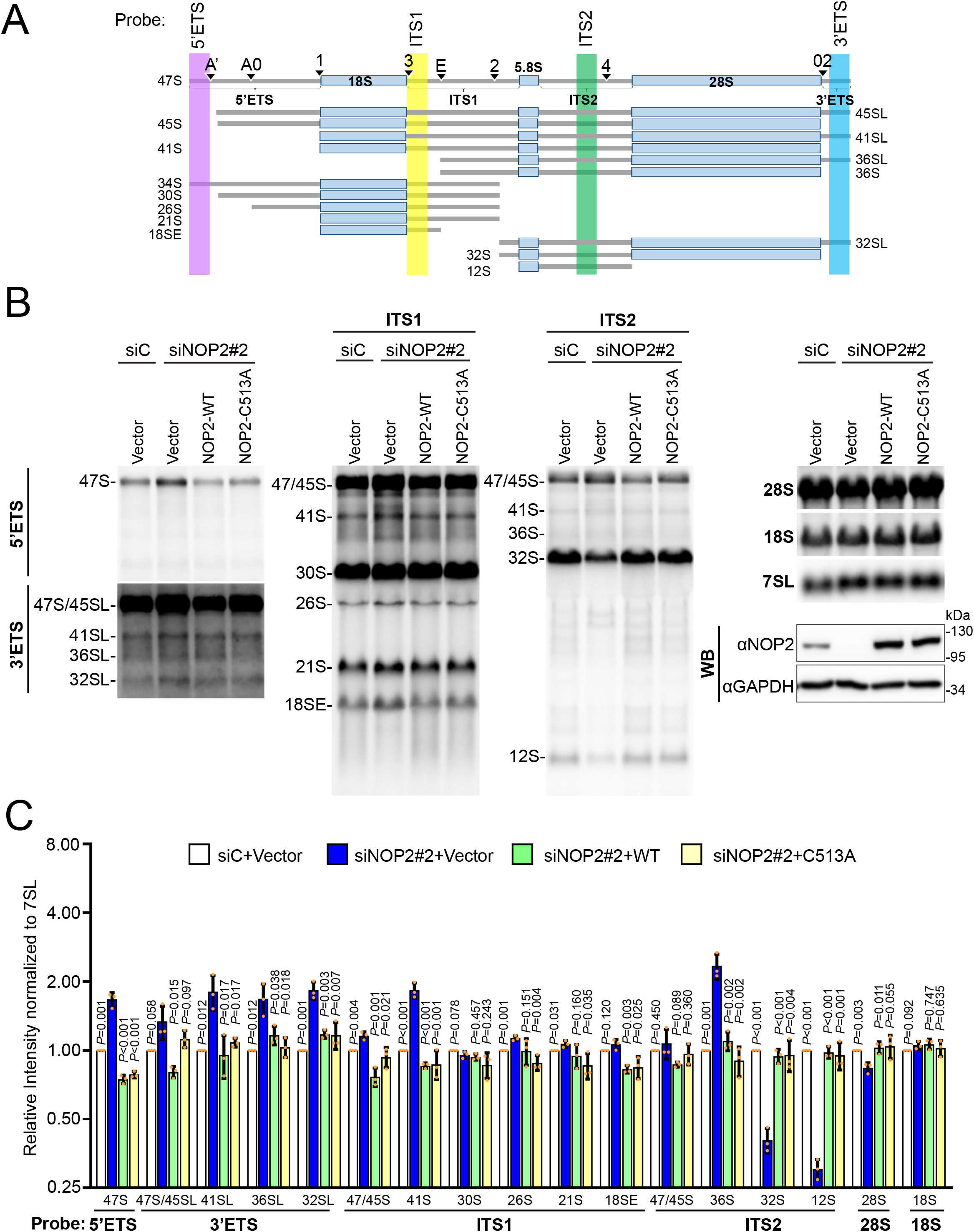
Human NOP2/NSUN1 is required for efficient pre-rRNA processing. **(A)** Schematic overview of human precursor rRNA processing and location of probes used for Northern blot indicated in colored areas (lavender, yellow, green and blue). **(B)** HCT116 cells expressing empty vector (Vector), siRNA resistant NOP2/NSUN1 WT, or the C513A catalytically inactive mutant were transfected with non-targeting control (siC) or NOP2 siRNA #2. After 72h, total RNA was separated on formaldehyde denaturing agarose gel and analyzed by Northern blot using 5’ETS, 3’ETS, ITS-1, ITS-2, 18S, 28S, and 7SL probes. A fraction of cells was collected to determine endogenous NOP2/NSUN1 depletion and ectopic expression of NOP2/NSUN1 WT or C513A by Western blot using a NOP2 antibody. **(C)** Densitometry quantification of each rRNA precursor from (B) normalized to 7SL RNA. The data are presented as the mean of 3 independent biological replicates ± standard deviation (SD). P-values comparing empty vector with NOP2/NSUN1 knockdown (siNOP2#2+Vector) were calculated using a 2-tailed independent student t-test.

To determine whether, NOP2/NSUN1 methyltransferase activity is required for rRNA processing, we performed rescue experiments by re-expressing siRNA resistant NOP2/NSUN1 WT or a catalytically inactive mutant (C513A) in NOP2/NSUN1 depleted cells. Proper knockdowns and ectopic re-expression of NOP2/NSUN1 were further confirmed by qRT-PCR (Supplementary Figure S5A). Remarkably, re-expression of both NOP2/NSUN1 WT and the C513A mutant was able to rescue all processing defects observed upon NOP2/NSUN1 depletion, including the decrease in 32S and 12S intermediates (Figure 5B-C and Supplementary Figure S4B showing different blot exposures). Bisulfite sequencing experiments on RNA from the knock-down cells re-expressing NOP2/NSUN1 WT or the C513A mutant confirmed the inactivity of the C513A mutant, which did not rescue the decrease of m^5^C4447 observed contrary to WT (Supplementary Figure S5B-D). Expression of the C513A mutant in the NOP2/NSUN1 RNAi background led to an increase of converted C4447 to 60%, possibly because this catalytically inactive mutant is competing with the function of the residual endogenous NSUN1/NOP2. Together these findings indicate that the methyltransferase catalytic activity of NOP2/NSUN1 is not required for rRNA processing.

Although steady-state levels of mature 28S rRNAs was only slightly reduced by NOP2/NSUN1 depletion (Supplementary Figure S3A-B) the decrease in the 32S and 12S intermediates observed suggest that NOP2/NSUN1, like its yeast homolog, is essential for 60S ribosomal subunit biogenesis (68). To confirm this finding, we performed polysome profiling to monitor the levels of 40S and 60S subunits, 80S monosomes and polysomes produced in the absence of NOP2/NSUN1. Because depletion of NOP2/NSUN1 by RNAi severely impairs cell proliferation (Figure 10C-D-F), we established stable cell lines expressing Dox-inducible NOP2/NSUN1 shRNA. We first verified that this depletion method by shRNA led to similar rRNA processing defects like the ones observed by acute RNAi depletion. As shown in Supplementary Figure S6A-B, comparable processing defects were detected upon shRNA depletion of NOP2/NSUN1, except for the increase in the 47S and 47/46S precursors, which is likely due to the more modest knockdown obtained with this method. Nevertheless, compared with uninduced cells, cells depleted of NOP2/NSUN1 by doxycycline treatment showed specific reduction in 60S levels as evidenced by the increase in 40S to 60S ratio and consequent decline in 80S monosomes (Figure 6A-B). While the levels of polysomes were not reduced by NOP2/NSUN1 depletion, puromycin labeling revealed a significant decrease in the overall translation rate (Figure 6C-D).

**Figure 6.**
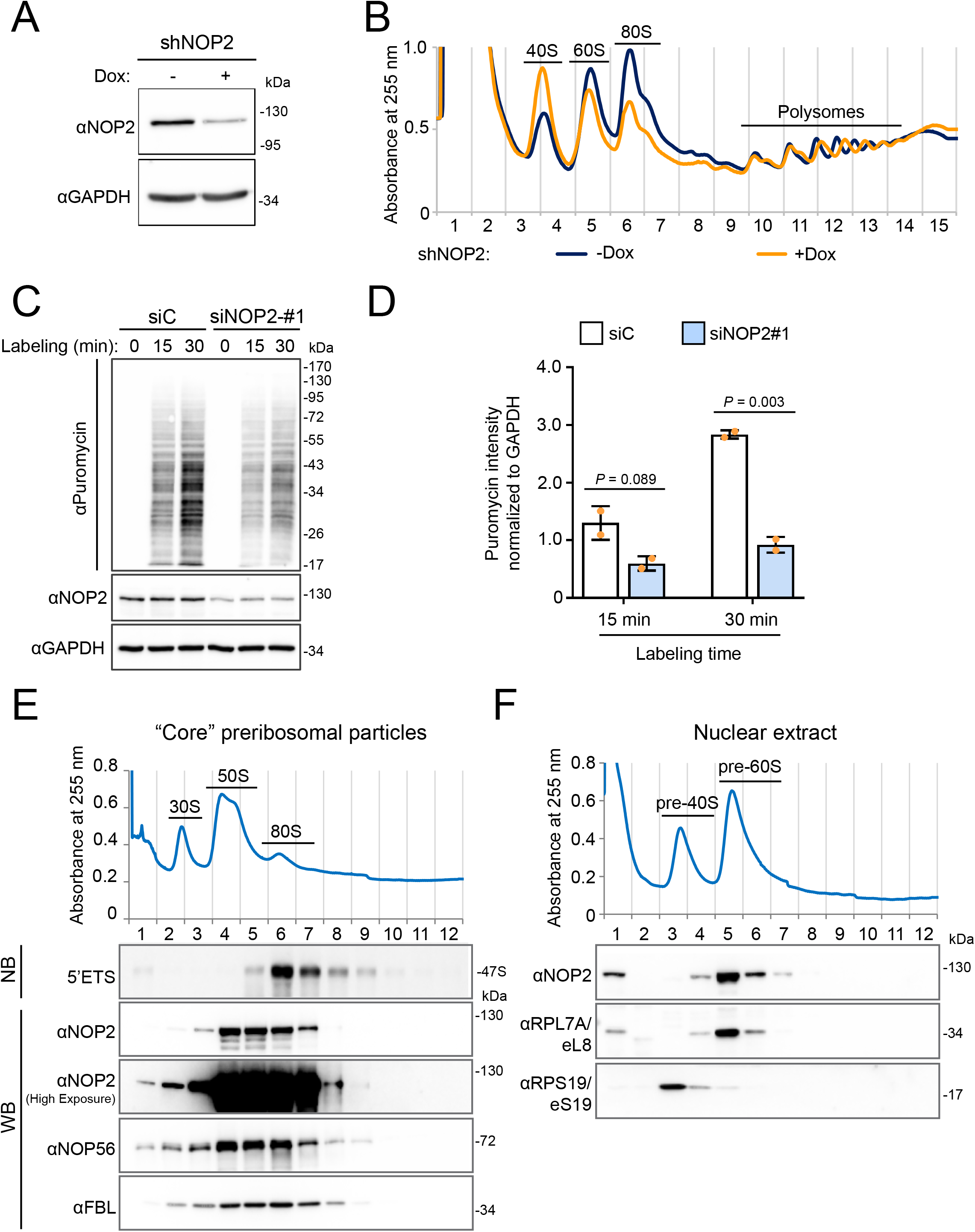
Human NOP2/NUSN1 is required for 60S ribosomal subunit biogenesis. **(A)** HCT116 cells were infected with doxycycline-inducible NOP2 shRNA expressing lentivirus. After puromycin selection, cells were induced with 200 ng/ml doxycycline (Dox+) for four days. Non-induced (Dox-) cells were used as control. NOP2/NSUN1 depletion efficiency was determined by Western blot. **(B)** A fraction of cells from (A) was analyzed by polysome profiling using total cell lysates. Dox+ and Dox-indicate doxycycline-induced and non-induced control, respectively. **(C)** HCT116 cells were transfected with non-targeting control (siC) or NOP2 siRNA. 72h after transfection, cells were treated with 5 µg/ml puromycin for the indicated time. Puromycylation of nascent peptides and NOP2/NSUN1 depletion efficiency, was determined by Western blot using puromycin and NOP2 antibodies. **(D)** Densitometry quantification of the puromycin signal from each lane from (C) was normalized to the corresponding GAPDH signal. The data is presented as the mean of 2 independent biological replicates ± standard deviation (SD). Statistical significance between NOP2 depleted samples and non-targeting control samples was calculated using a 2-tailed independent student *t*-test. **(E)** High molecular weight complexes containing pre-rRNAs and tightly associated ribosome assembly factors, referred to as “core” preribosomal particles isolated from HCT116 nuclear extracts under high-salt conditions were separated on sucrose gradient ultra-centrifugation followed by fractionation. RNA and proteins were isolated from each fraction and analyzed by Western blotting (WB) or Northern blot (NB) with the indicated antibodies or probe, respectively. **(F)** Nuclear extract from HCT116 cells was separated on sucrose gradient ultra-centrifugation followed by fractionation. Proteins were extracted from each fraction and analyzed by Western blot with the indicated antibodies.

To verify our miCLIP-seq data showing tight binding of NOP2/NSUN1 to the 5’ETS region of the pre-rRNA, we performed sucrose gradient fractionations of nuclear pre-ribosomes followed by Western blotting to determine the co-sedimentation pattern of NOP2/NSUN1. High molecular weight complexes containing pre-rRNAs and tightly associated ribosome assembly factors were isolated as “core” pre-ribosomal particles under high-salt conditions (69). These conditions are known to resolve three major peaks at around 30S, 50S and 80S (49,69,70). Figure 6E shows that NOP2/NSUN1 mainly co-sediments with the 50S and 80S peaks, which were previously shown to represent core particles respectively containing the 32S rRNA intermediate and the 47S pre-rRNA (Figure 6E 5’ETS Northern blot panel) (49, 70). Sucrose gradient fractionation of nuclear extract isolated in milder conditions showed that NOP2/NSUN1 co-sediments with the pre-60S particles (Figure 6F), corroborating the co-sedimentation with the 32S rRNA precursor from the 50S core pre-ribosomes (Figure 6E). Together, these findings indicate that NOP2/NSUN1 tightly associates with particles containing the 47S pre-rRNA as well as pre-60S ribosomes, which is consistent with a role for NOP2/NSUN1 in regulating U3-dependent 5’ETS processing likely in the 90S complex, as well as U8-dependent cleavage reactions leading to synthesis of the large subunit rRNAs (pre-60S particle).

### NOP2/NSUN1 regulates snoRNP functions

Our CLIP-seq analysis revealed that NOP2/NSUN1 likely interacts in a non-catalytic manner with box C/D snoRNAs, which we independently confirmed by NOP2/NSUN1 immunoprecipitation combined with qRT-PCR analysis (Supplementary Figure S1B). The rRNA processing defects we observed after NOP2/NSUN1 depletion are very similar to those observed after depletion of U3 and U8, suggesting that NOP2/NSUN1 may be important for regulating their rRNA processing-related functions. U3 and U8 also contain the conserved box C/D motif, which is recognized by 15.5K, NOP56, NOP58, and Fibrillarin; the four core proteins together with the snoRNAs that assemble to form rRNA processing snoRNPs (46). To investigate the roles of NOP2/NSUN1 in box C/D snoRNAs regulation, we first tested whether NOP2/NSUN1 associates with the core proteins NOP56 and Fibrillarin by co-immunoprecipitation experiments. All commercially available NOP2/NSUN1 antibodies tested could not precipitate any detectable NOP2/NSUN1 proteins so we used a FLAG-tagged version to perform the interaction assay. As shown in Figure 7A, ectopically expressed FLAG-NOP2/NSUN1 co-precipitated with both NOP56 and Fibrillarin. Reversely, immunoprecipitation of NOP56 also co-precipitated endogenous NOP2/NSUN1, indicating that these proteins form a complex at physiological level (Figure 7B). We next tested whether NOP2/NSUN1 is required for snoRNPs complex stability. Depletion of NOP2/NSUN1 by RNAi did not affect NOP56 and Fibrillarin protein levels nor the ability of NOP56 to interact with Fibrillarin as detected by co-immunoprecipitation and Western blotting experiments (Figure 7C and Supplementary Figure S7E shows the densitometry quantification). To determine whether NOP2/NSUN1 is required for nucleolar snoRNP complex stability we immuno-precipitated NOP56 or 15.5K and tested the presence of box C/D snoRNAs, including U3 and U8, in the complex. A fraction of the immuno-precipitation was kept for Western blot analysis to monitor the immuno-precipitation efficiency (Supplementary Figure S7A and C) and the rest was processed for RNA extraction and qRT-PCR analysis (Supplementary Figure S7B and D). We observed a marked decrease in the level of snoRNAs bound to NOP56 or 15.5K after knockdown of NOP2/NSUN1 (Supplementary Figure S7A-D). Steady-state levels of U3 and U8 snoRNAs were not affected by NOP2/NSUN1 depletion (Supplementary Figure S4C-D), suggesting that NOP2/NSUN1 is likely required for some aspects of nucleolar snoRNPs complex stability. As predicted by our miCLIP-seq data, the catalytic activity of NOP2/NSUN1 is not necessary for the regulation of snoRNPs complex stability as re-expression of both NOP2/NSUN1 WT and the C513A mutant rescued the decrease in snoRNAs bound to NOP56 or 15.5K after NOP2/NSUN1 depletion (Figure 7D-G).

**Figure 7.**
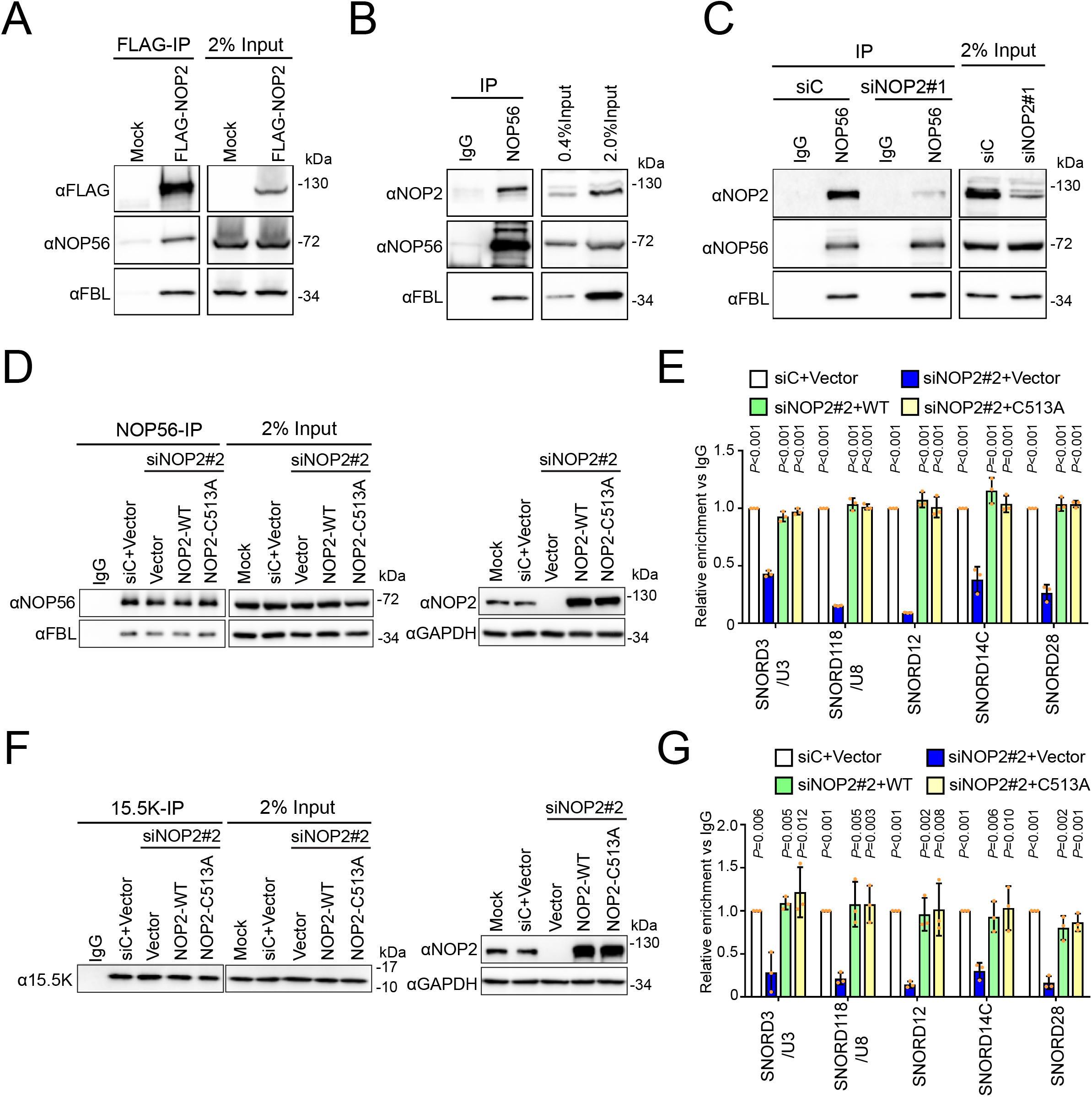
NOP2/NSUN1 is required to maintain the integrity of C/D box snoRNPs. **(A)** HEK293T expressing empty vector (Mock) or FLAG-tagged NOP2/NUSN1 WT were lysed and immunoprecipitated (IP) with an anti-FLAG antibody. Associated proteins were analyzed by Western blot with the indicated antibodies. Mock transfected cells were used as negative control. **(B)** NOP56 was immunoprecipitated with an anti-NOP56 antibody from HEK293T cell lysate. Associated proteins were analyzed by Western blot with the indicated antibodies. Normal rabbit IgG was used as a negative immunoprecipitation control. **(C)** HEK293T cells were transfected with non-targeting control (siC) or NOP2 siRNA. 72h later, cells were lysed and immunoprecipitated (IP) with an anti-NOP56 antibody. Associated proteins were analyzed by Western blot with the indicated antibodies. Normal rabbit IgG was used as a negative immunoprecipitation control. **(D-G)** HCT116 cells expressing empty vector (Vector), siRNA resistant NOP2/NSUN1 WT or the C513A catalytically inactive mutant were transfected with non-targeting control (siC) or NOP2 siRNA #2. After 72h, cells were lysed and immunoprecipitated (IP) with anti-NOP56 **(D, E)** or anti-15.5K **(F, G)** antibody. A fraction of associated proteins was analyzed by Western blot to control for immunoprecipitation and knockdown efficiency **(D, F)**. The remaining immunoprecipitation fraction was processed for RNA extraction followed by RT-qPCR using SNORD3/U3, SNORD118/U8, SNORD12, SNORD14C, and SNORD28 specific primers **(E, G)**. Relative enrichment over IgG control is represented. Data are presented as mean of 3 independent biological replicates ± standard deviation (SD). Statistical significance values relative to empty vector NOP2/NSUN1 knockdown (siNOP2+Vector) were calculated using a 2-tailed independent student t-test.

The biogenesis of box C/D snoRNPs occurs in the nucleoplasm through the chaperoning action of large non-snoRNP multiprotein complexes involved in core protein assembly and maturation of the snoRNA (71). Previous studies have shown that the biogenesis process of U3 and U8 snoRNPs involves restructuring events that stabilizes the association of the core box C/D proteins to the snoRNA just before or during their localization to the nucleolus (46, 72). To determine if NOP2/NSUN1 plays a role in box C/D snoRNPs biogenesis, we first tested whether NOP2/NSUN1 interacts with U3 and U8 snoRNPs in the nucleoplasm or nucleolus. Nuclear and nucleolar extracts were prepared from FLAG-tagged NOP2/NSUN1 WT expressing cells and the association of NOP2/NSUN1 with NOP56 and Fibrillarin snoRNP proteins and U3 and U8 snoRNA was assessed by immunoprecipitating NOP2/NSUN1 from each compartment with a FLAG antibody followed by Western blotting or qRT-PCR analysis, respectively. Both nuclear and nucleolar NOP2/NSUN1 associated with the core snoRNP proteins NOP56 and Fibrillarin (Figure 8A) as well as with U3 and U8 snoRNAs (Figure 8B). Reciprocally, immunoprecipitation of endogenous NOP56 shows that NOP2/NSUN1 interacts with snoRNPs in the nucleus and nucleolus (Figure 8C). The input fractions in Figure 8C show an underrepresented nucleolar content as we could not completely extract it by sonication without causing lysate precipitation. Immunofluorescence experiment and biochemical analysis of NOP2/NSUN1 subcellular localization demonstrate that NOP2/NSUN1 localizes majorly to the nucleolus with a smaller proportion observed in the nucleus (Supplementary Figure S8A-B). Nevertheless, our data indicate that NOP2/NSUN1 associates with box C/D snoRNPs in the nucleus but that most of the interaction occurs in the nucleolar compartment. Together, these findings suggest that NOP2/NSUN1 may regulate both the biogenesis of box C/D snoRNPs in the nucleus and their recruitment to pre-ribosomes in the nucleolus. In line with this, we find that NOP2/NSUN1 interacts with the box C/D snoRNPs assembly factors NUFIP1 and RUVBL1 by co-immunoprecipitation experiments at physiological levels (Figure 8D). The association of NOP2/NSUN1 with NUFIP1 has previously been reported in large scale analysis of the human interactome, solidifying our findings (73). NOP2/NSUN1 did not co-precipitate with La protein, a biogenesis factor that associates with U3 snoRNA precursor and that is predicted to have a role in U3 3’-end processing (Figure 8E). Although we did not test all known biogenesis factors, our data demonstrate a specificity of interaction for NSUN1/NOP2 with biogenesis factors involved in snoRNP assembly.

**Figure 8.**
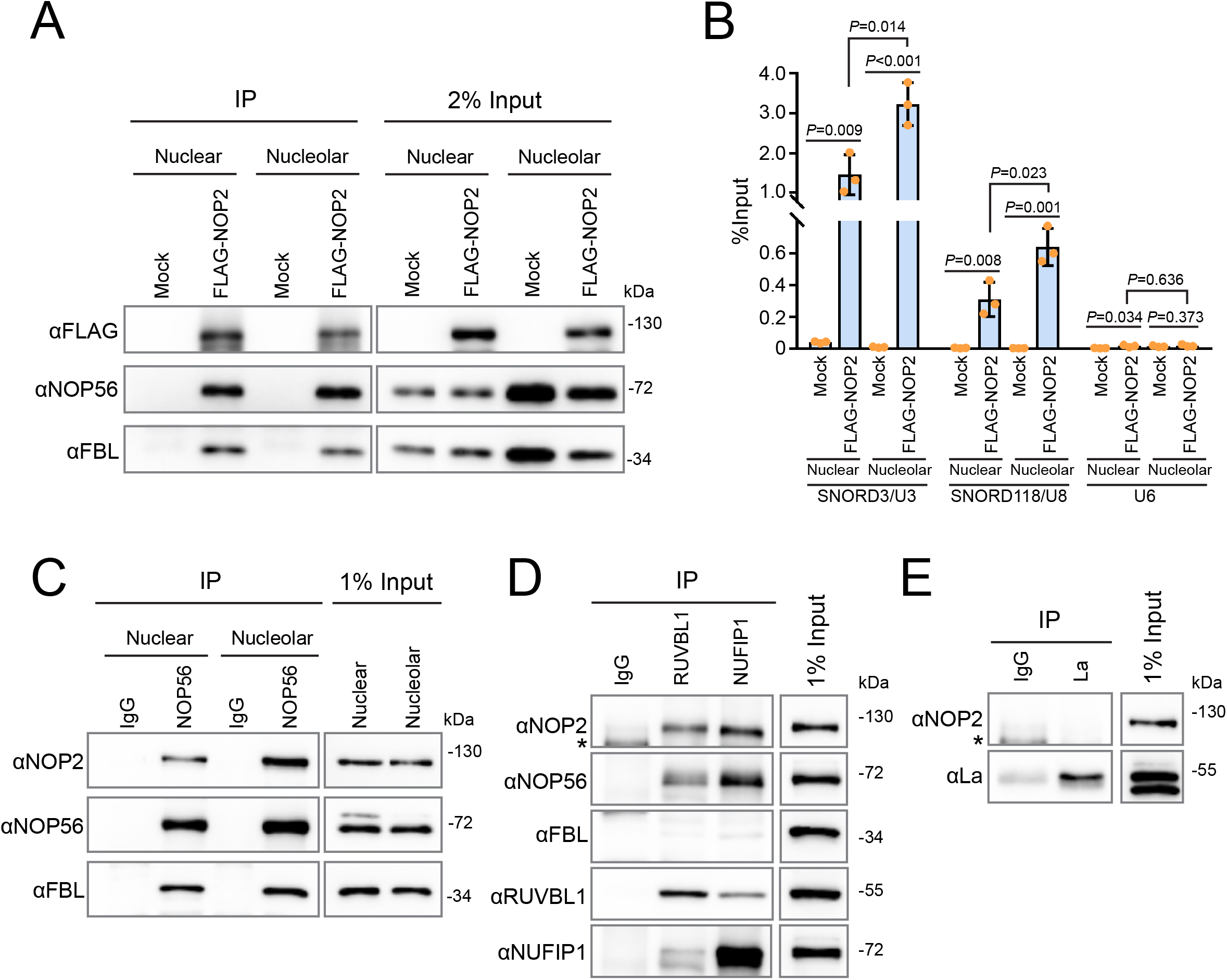
NOP2/NSUN1 interacts with nuclear and nucleolar snoRNPs. **(A)** Nuclear and nucleolar extracts from HEK293T cells expressing empty vector (Mock) or FLAG-tagged NOP2/NUSN1 WT were immunoprecipitated (IP) with an anti-FLAG antibody. A fraction of the IP sample was analyzed by Western blot. **(B)** RNA from the remaining IP sample in (A) was extracted and analyzed by RT-qPCR with SNORD3/U3, SNORD118/U8, and U6 specific primers. **(C)** Nuclear and nucleolar extracts from HCT116 cells were immunoprecipitated with an anti-NOP56 antibody. Associated proteins were analyzed by Western blot. Normal rabbit IgG was used as negative immunoprecipitation control. **(D, E)** HCT116 cells were lysed and immunoprecipitated with anti-RUVBL1, anti-NUFIP1 **(D)**, or anti-La **(E)** antibodies. Associated proteins were analyzed by Western blot. Normal rabbit IgG was used as negative immunoprecipitation control. Asterisk (*) indicates non-specific bands.

As our data show that NOP2/NSUN1 co-sediments with the 47S pre-rRNA and that it binds to both the 5’ETS pre-rRNA region and to box C/D snoRNAs in the nucleolus, it suggests that NOP2/NSUN1 may also be important for recruiting box C/D snoRNPs to the 90S particle. To assess this, we performed sucrose gradient fractionations of isolated “core” pre-ribosomes followed by Northern blotting to monitor changes in the co-sedimentation pattern of U3 and U8 after NOP2/NSUN1 depletion. In agreement with previous studies (69, 74), U8 snoRNA fractionated in two gradient peaks (fraction 1 and fraction 6 (80S)), while U3 fractionated in three peaks (fractions 1, 4+5 (50S) and 6 (80S)) (Figure 9A-C). From our Northern blot analysis (Supplementary Figure S9) and previous reports (69), fraction 1 represents free snoRNPs, fractions 4+5 represent pre-ribosomes with ITS1 and ITS2-containing rRNA intermediates, and fraction 6 represents early 47S particles. Compared with uninduced cells, cells depleted of NOP2/NSUN1 by doxycycline-induced shRNA knockdown showed a modest but reproducible reduction in the levels of both U3 and U8 co-sedimenting with the 47S pre-rRNA (Figure 9A-C). Notably, the distribution of U8 and U3 broadened over to fractions 4 and 5 in NOP2/NSUN1 depleted cells, indicating a persistence in pre-40S or late pre-60S particles. The reason for this is not apparent and will require further investigation. It is possible that this is due to a secondary effect caused by the slowed rate of early pre-rRNA processing or the release of U8 and U3 snoRNPs from pre-ribosomal particles is impaired in the absence of NOP2/NSUN1.

**Figure 9.**
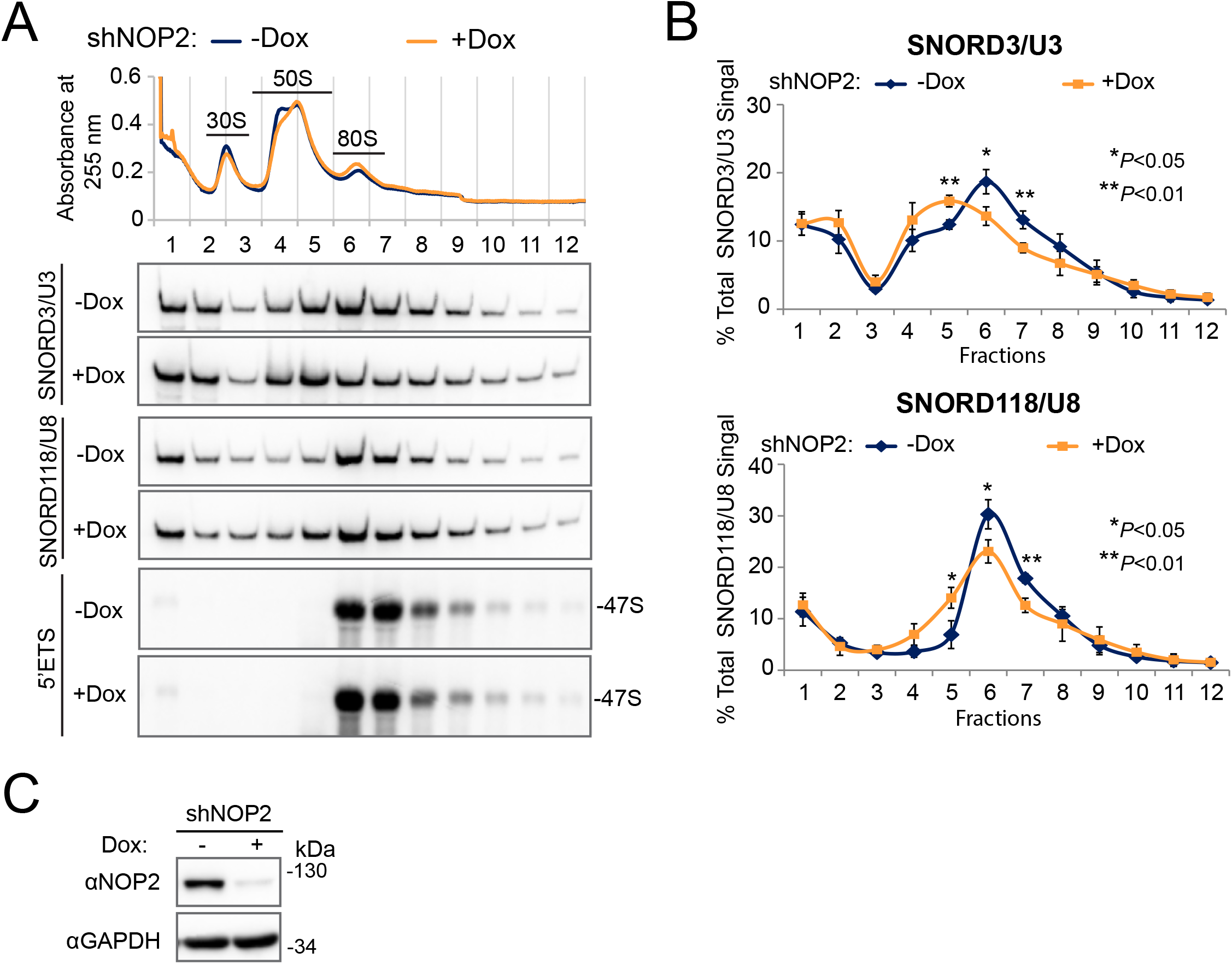
Depletion of NOP2/NSUN1 reduces SNORD3/U3 and SNORD118/U8 co-sedimentation with 47S rRNA-containing pre-ribosomal particles. **(A)** HCT116 cells expressing doxycycline-inducible NOP2 shRNA were induced with 200 ng/ml doxycycline (Dox+) for 6 days. Non-induced (Dox-) cells were used as control. High molecular weight complexes containing pre-rRNAs and tightly associated ribosome assembly factors isolated from HCT116 nuclear extracts under high-salt conditions were separated on sucrose gradient ultra-centrifugation followed by fractionation. RNA was isolated from each fraction and analyzed by Northern blot using SNORD3/U3, SNORD118/U8 and 5’ETS probes. **(B)** Densitometry quantification of SNORD3/U3 and SNORD118/U8 signal from (A) normalized to the respective total signal from all fractions. Data are presented as the mean of 3 independent biological replicates ± standard deviation (SD). Statistical significance between NOP2 depleted samples (Dox+) and non-induced control samples (Dox-) was calculated using a 2-tailed independent student t-test. **(C)** A small fraction of cells from (A) were collected to determine NOP2/NSUN1 depletion efficiency using Western blot with the indicated antibodies.

Inhibition of any steps of ribosome biogenesis is known to induce a nucleolar stress characterized by activation of p53 and subsequent cell cycle arrest (75–88). As previously reported, depletion of NOP2/NSUN1 by RNAi caused inhibition of cell proliferation and induced ribosomal stress as evidenced by stabilization of p53 and activation of its transcriptional target p21^CIP1^ (Figure 10A-C) (47, 89). However, we find that NOP2/NSUN1 knockdown also impaired cell proliferation (Figure 10D) and compromised 5’ETS maturation and levels of 32S and 12S intermediates in p53-null cells (HCT116 p53 -/-) (Supplementary Figure S10). Remarkably, re-expression of NOP2/NSUN1 WT or the catalytically inactive mutant C513A in the NOP2/NSUN1 knockdown background were both able to rescue the impaired cell proliferation and the activation of p53 (Figure 10E-F). This suggests that the rRNA processing defects observed are likely due to a loss of NOP2/NSUN1-dependent regulation of snoRNPs complex formation and not caused by loss of 28S rRNA C4447 methylation or by p53 activation. Taken together, our findings indicate that the m^5^C catalytic activity of NOP2/NSUN1 is not critical for rRNA processing and cell proliferation.

**Figure 10.**
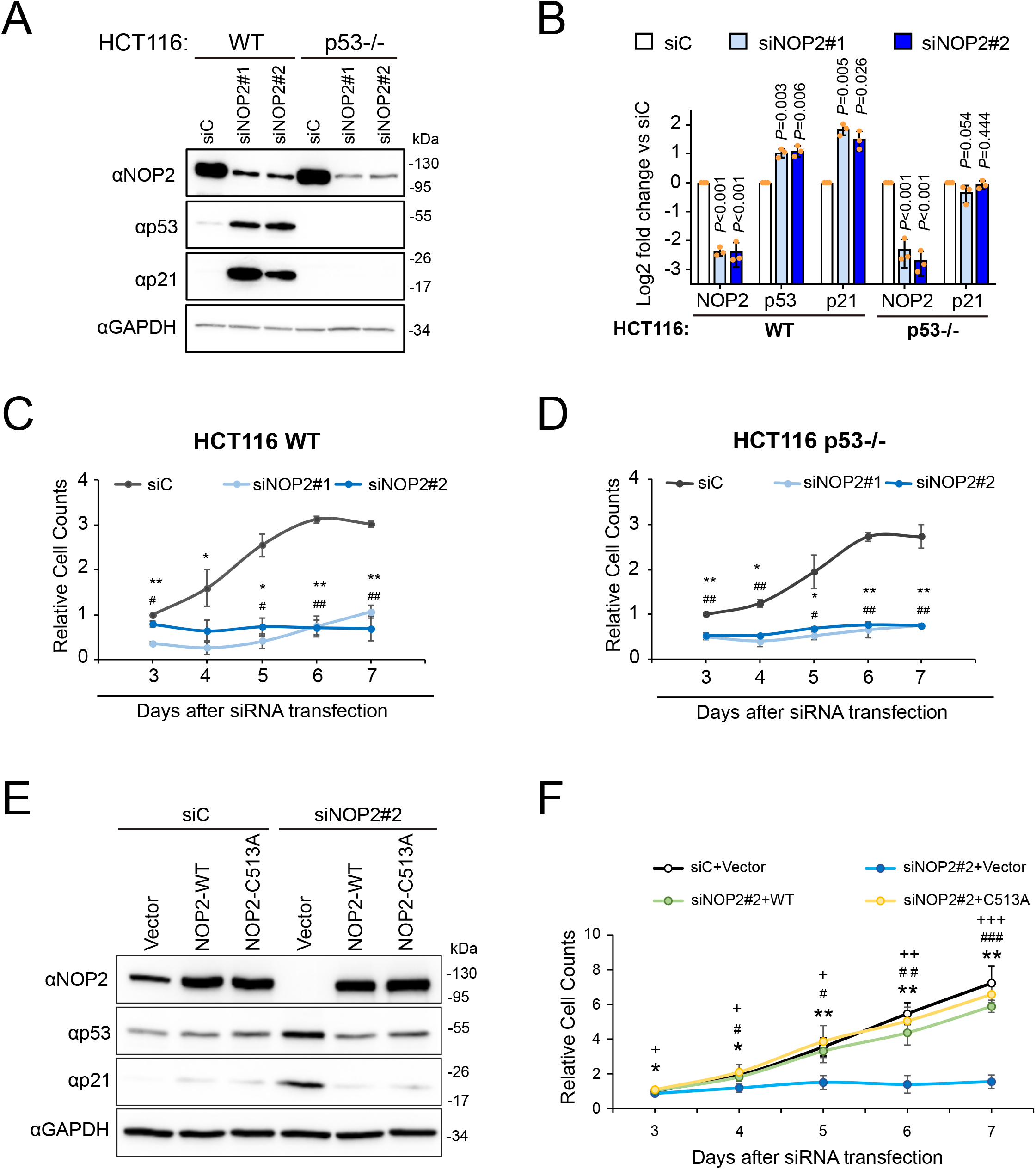
The RNA methyltransferase activity of NOP2/NSUN1 is not required to maintain cell proliferation. **(A)** HCT116 WT or p53 -/- cells were transfected with non-targeting control (siC), NOP2 siRNA #1 or #2. 72h later, a fraction of the cells was harvested for Western blot analysis with specific antibodies as indicated. **(B)** Densitometry quantification of each protein from (A) normalized to GAPDH. Data are presented as the mean of 3 independent biological replicates ± standard deviation (SD). Statistical significance between NOP2 depleted samples and non-targeting control samples was calculated using a 2-tailed independent student *t*-test. A fraction of WT **(C)** and p53 -/- **(D)** HCT116 cells from (A) were monitored for proliferation daily using an MTT assay. Relative cell counts were normalized to non-targeting control (siC) on day 3. Data are presented as the mean of 3 independent biological replicates ± standard deviation (SD). Statistical significance between NOP2 depleted samples and non-targeting control samples was calculated using a 2-tailed independent student *t*-test. *P<0.05 and **P<0.01: NOP2 siRNA#1 vs siC; ^#^P<0.05 and ^##^P<0.01: NOP2 siRNA#2 vs siC. **(E)** HCT116 cells expressing empty vector (Vector), siRNA resistant NOP2/NSUN1 WT or C513A mutant were transfected with non-targeting control (siC) or NOP2 siRNA #2. After 72h, a fraction of cells was collected for Western blot analysis with specific antibodies as indicated. **(F)** A small fraction of cells from (E) was monitored daily for proliferation using an MTT assay. Relative cell counts were normalized to empty vector with non-targeting control (siC+Vector) on day 3. Data are presented as the mean of 3 independent biological replicates ± standard deviation (SD). Statistical significance values relative to empty vector + NOP2/NSUN1 knockdown (siNOP2#2+Vector) were calculated using 2-tailed independent student t-test. *P<0.05 and **P<0.01: vs siC; ^#^P<0.05, ^##^P<0.01 and ^###^P<0.001: vs siNOP2#2+WT; ^+^P<0.05, ^++^P<0.01 and ^+++^P<0.001: vs siNOP2#2+C513A.

## DISCUSSION

In this study we used a genome-wide approach to identify RNA methylation targets of human NOP2/NSUN1 to define its role in ribosome biogenesis. We adapted a miCLIP approach using a catalytic mutant of NOP2/NSUN1 that allowed the purification of covalently cross-linked NOP2/NSUN1 to its methylated cytosine on substrate RNA. Our findings reveal multiple roles played by NOP2/NSUN1 in the regulation of ribosome biogenesis. One function involves the deposition of m^5^C at position C4447 of the 28S rRNA, which has been previously hypothesized to help stabilize rRNA structure and contribute to efficient ITS2 processing and large subunit assembly. However, our findings reveal that NOP2/NSUN1 catalytic activity and the deposition of m^5^C on the 28S rRNA are not critical for efficient rRNA processing and to maintain cell proliferation. Unexpectedly, our study also uncovered a function for NOP2/NSUN1 in box C/D snoRNPs regulation independent of its catalytic activity, and most likely explains the complex rRNA processing defects resulting from NOP2/NSUN1 depletion.

Several studies have suggested that NOP2/NSUN1 performs additional functions independent of its C5 cytosine methylation activity (13, 37). In budding yeast, the lethality of the NOP2/NSUN1-deleted strain can be rescued by re-expression of a methyltransferase-dead mutant, indicating the catalytic function of NOP2/NSUN1 is not essential for viability (37). In like manner, knockout of NOP2/NSUN1 is lethal in worms whereas mutant worms expressing a catalytically-dead NOP2/NSUN1 remain viable (13). Our findings suggest that box C/D snoRNPs regulation is an essential non-catalytic function of NOP2/NSUN1 that likely contributes to the lethality phenotype of NOP2/NSUN1 depletion. Our data is consistent with previous findings showing that other rRNA modifying enzymes, such as EMG1, Dim1p and WBSCR22 (Bud23), bind to the pre-rRNA at an early stage to coordinate processing assembly events independently of their catalytic functions (90–93).

In contrast to budding yeast, in which the 35S and 27S pre-rRNA transcripts accumulate in the absence of NOP2/NSUN1 (68), depletion of NOP2/NSUN1 in human cells leads to a more severe rRNA processing defect marked by the accumulation of the 47S primary transcript and a distinct reduction of both 32S and 12S rRNA precursors (Figure 5 and Supplementary Figure S3). Processing of ITS1 and 3’ETS regions was also affected. These rRNA processing defects are highly similar to those observed with depletion of the U3 and U8 box C/D snoRNAs (65). Indeed, we found that the recruitment of U3 and U8 to pre-ribosomes (Figure 9) and their stable complex formation with NOP56 and 15.5K was compromised in the absence of NOP2/NSUN1 (Figure 7D-G and Supplementary Figure S7A-D). In addition, our data shows that NOP2/NSUN1 interacts with snoRNP biogenesis factors and with snoRNPs from nucleoplasm and nucleolar fractions (Figure 8). These observations, in addition to our finding that NOP2/NSUN1 binds to the rRNA 5’ETS, suggest that NOP2/NSUN1 is important for U3 and U8 snoRNP complexes assembly in the nucleus and possibly their recruitment to the 90S particle in the nucleolus. Interestingly, most box C/D snoRNP biogenesis factors are essential to maintaining box C/D snoRNA levels (46, 94). However, depletion of NOP2/NSUN1 did not lead to detectable changes in U3 or U8 steady-state levels (Supplementary Figure S4C-D). Further work will be required to determine the exact role of NOP2/NSUN1 in snoRNP assembly.

NOP2/NSUN1 is a component of the human UTP-B complex, a stable subcomplex of 90S assembly factors, that also includes UTP-A, UTP-C and U3 snoRNP (95, 96). Although its functions are not entirely understood in humans, yeast UTP-B is likely acting as an RNA chaperone critical to initiating the assembly of the 90S pre-ribosomal particle and depletion of some of its components has been shown to impair U3 snoRNP recruitment and processing of the pre-rRNA 5’ETS (97–101). Our CLIP analysis revealed that NOP2/NSUN1 binds to the 5’ETS region and U3 snoRNA (Figure 3A-B and Supplementary Figure S2C) and that its depletion reduces the amount of U3 present in particles containing the 47S pre-rRNA (Figure 9). These findings agree with a role for NOP2/NSUN1 as a UTP-B complex component and may also explain the accumulation of unprocessed 47S/45S primary transcript observed upon NOP2/NSUN1 knockdown. Although we found that NOP2/NSUN1 binds broadly over the entire 5’ETS with some peaks overlapping with U3 snoRNA 5’ hinge binding site, most CLIP peaks were enriched between the A0 and 1 cleavage sites of the 5’ETS (101) (Figure 3A). In contrast, other UTP-B factors such as UTP1/PWP2, UTP12/WDR3 and UTP18 have been shown to make contact on the 5’ETS upstream of the A0 site (50, 101). As U3 snoRNA-5′ETS base-pairing is thought to be facilitated by the UTP-B complex (97,98,101), it is possible that NOP2/NSUN1 could also have a dual role in stabilizing the 5′ ETS regions downstream of the A0 site while facilitating the recruitment of U3 snoRNP and UTP-B for effective rRNA processing and ribosome assembly. It remains to be determined whether NOP2/NSUN1 is required for the assembly of the UTP-B complex or its recruitment to the pre-90S particle.

U8 is a vertebrate-specific snoRNA that was shown to be required for rRNA processing events leading to the synthesis of the 5.8S and 28S rRNAs in Xenopus (102, 103), mouse (104) and human (65) cells. Based on sequence homology and accessibility measurements, previous studies suggested that U8 snoRNA base pair with the 5’end of the 28S rRNA (see mapped position in Figure 3A-B) (103). Consequently, it has been proposed that U8 snoRNPs chaperone the formation of an ITS2 proximal stem required for proper processing (104). In yeast, ITS2 has been modeled to form distinct secondary structures that could independently facilitate processing events in the absence of U8 (105). The difference in U8 requirement for ITS2 maturation in vertebrates may explain the more severe processing defects we observed with NOP2/NSUN1 depletion. As our data indicate that NOP2/NSUN1 is required for the stability of U8 snoRNP complexes and the recruitment of U8 to pre-ribosomes (Figures 7 and 9), it is possible that the decline in 32S and 12S pre-rRNAs observed with NOP2/NSUN1 or U8 knockdown (65) is due to improperly folded ITS2-containing precursors that could be processed for degradation. Intriguingly, RNA duplex mapping in living cells uncovered that U8 mainly interacts with the 3’end of the 28S rRNA in a region that in close structural proximity to NOP2/NSUN1 methylation site at C4447 (see mapped position in Figure 3A-B) (106). The role of this interaction remains unclear and further work is needed to determine whether U8 is required to facilitate NOP2/NSUN1 access to its target site.

Recent studies in *C. elegans* have suggested that NOP2/NSUN1 is responsible for catalyzing the deposition of m^5^C at C2982 of the 26S rRNA (the equivalent of C4447 on human 28S rRNA) (42). Soma-specific RNAi depletion of NOP2/NSUN1 reduced worm body size, impaired fecundity, and gonad maturation while extending lifespan. Surprisingly, rRNA processing, 60S ribosome biogenesis and global protein synthesis were not found to be impaired by NOP2/NSUN1 depletion. Instead, changes in the translation pattern of specific mRNAs were observed, suggesting that NOP2/NSUN1 could play a role in modulating ribosome function (42). Our miCLIP analysis revealed that NOP2/NSUN1 binds to several box C/D snoRNAs (Supplementary Table 2) involved in guiding Fibrillarin-catalyzed 2’-O-methylation on rRNAs. We are investigating whether depletion of NOP2/NSUN1 impairs rRNA 2’-O-methylation, which would likely affect ribosome function and explain the changes in mRNA translation patterns observed in worms. In contrast to the phenotypes observed in worms, our data demonstrate that human NOP2/NSUN1 is essential for proliferation, rRNA processing and ribosome biogenesis. In line with an essential role for NOP2/NSUN1, our attempts to generate CRISPR knockouts were not successful and the only clones that could be recovered were NOP2/NSUN1 mutant heterozygotes alleles or homozygous mutants with destabilized mRNA leading to low levels of NOP2/NSUN1 protein production (not shown). NOP2/NSUN1 is overexpressed in a wide variety of cancer, and its expression level correlates with tumor aggressiveness, poor prognosis and therapy resistance (22,23,25- 35,107). NOP2/NSUN1 depletion has been shown to suppress proliferation, migration, and invasion of colon cancer cells (107) and sensitize leukemia cells to 5-aza-cytidine treatment (108). Although inhibition of ribosome biogenesis is known to induce cell cycle arrest through p53 activation (75–88), our data demonstrate that depletion of NOP2/NSUN1 severely impairs cell proliferation in both WT and p53-null cells. Together, these observations suggest that targeted therapies interfering with NOP2/NSUN1 functions would benefit a broad range of tumor types, regardless of p53 status.

## DATA AVAILABILITY

The datasets generated and analyzed during the current study are publicly available at the GEO genomics data repository under accession number GSE188735.

## SUPPLEMENTARY DATA

Supplementary Data are available at NAR Online.

---

**Supplementary Figure S1.**
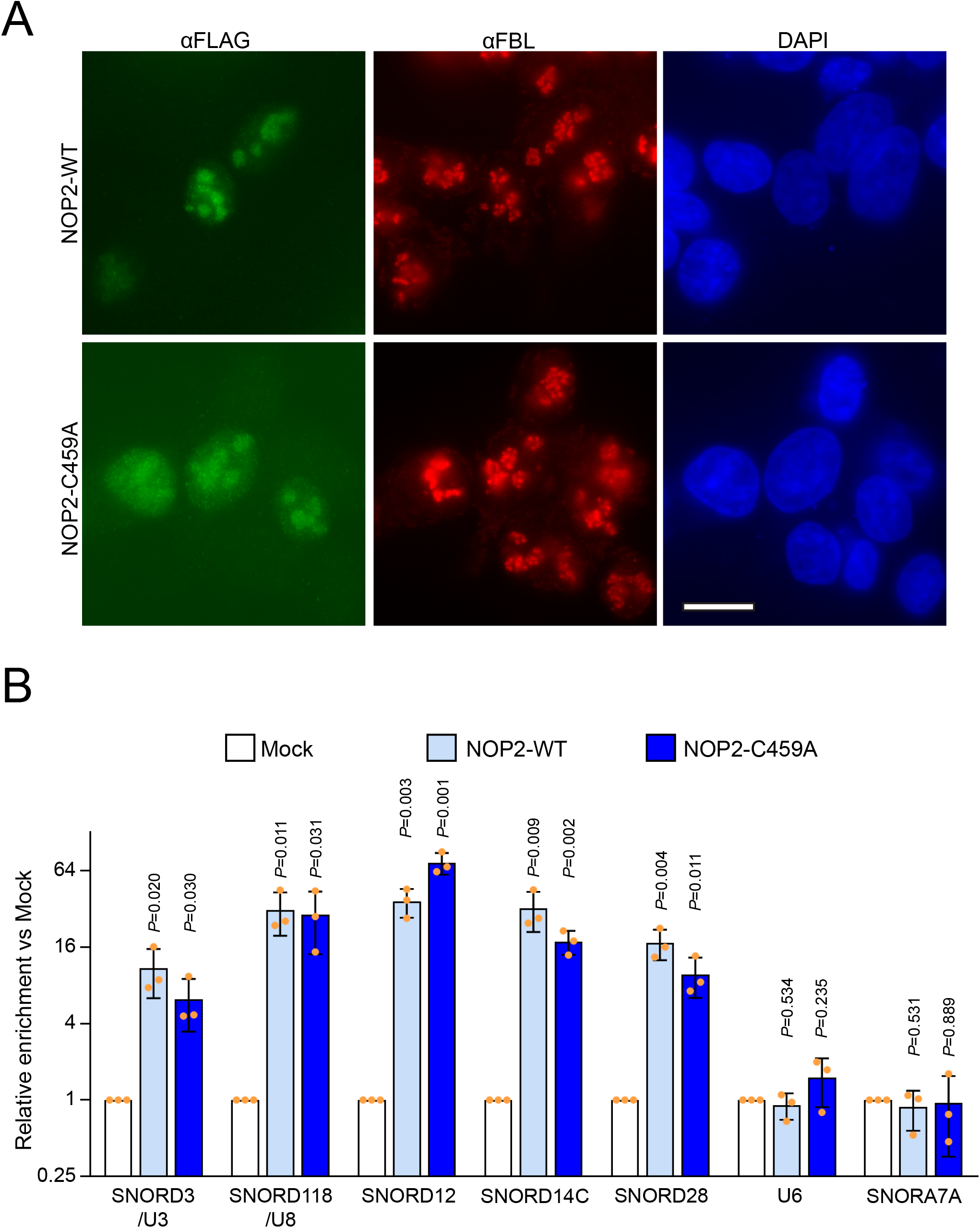
NOP2/NSUN1 binds to C/D box snoRNAs. **(A)** HEK293T cells expressing FLAG-tagged NOP2/NSUN1 WT or C459A mutant were fixed and immunostained with an anti-FLAG (green) and anti-FBL (red, nucleolar marker). DNA was visualized by staining with Hoechst 33342 (blue). The scale bar is representative of all panels: 10 μm. **(B)** HEK293T cells expressing FLAG-tagged NOP2/NSUN1 WT or C459A mutant were lysed and immunoprecipitated (IP) with an anti-FLAG antibody. Associated RNAs were analyzed by RT-qPCR with specific primers as indicated. Mock transfected cells immunoprecipitated (IP) with an anti-FLAG antibody were used as negative control. Enrichment was calculated over IgG control. Data are presented as the mean of 3 independent biological replicates ± standard deviation (SD). Statistical significance between Mock and WT or C459A mutant IP samples was calculated using a 2-tailed independent student *t*-test.

**Supplementary Figure S2.**
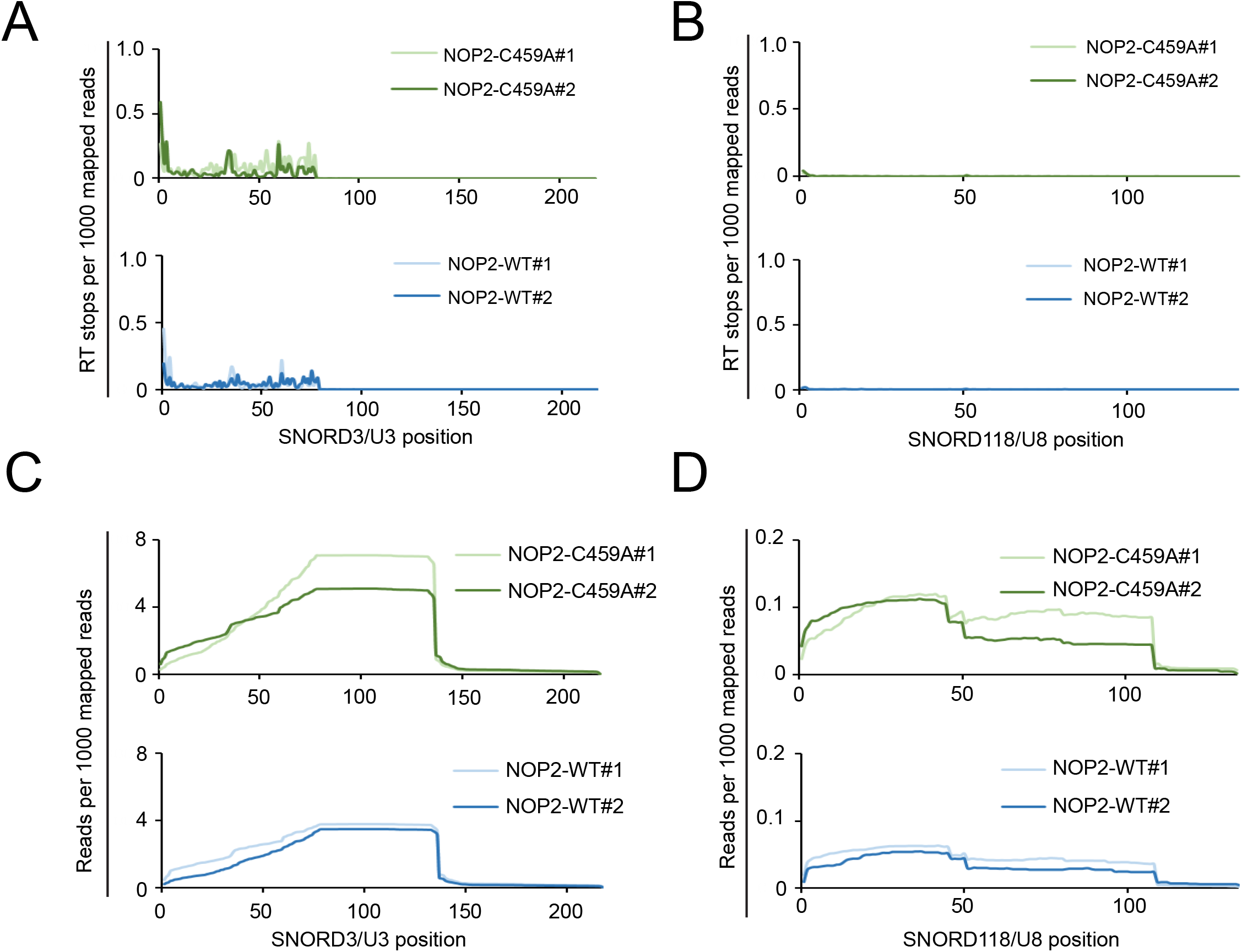
NOP2/NSUN1 does not crosslink to C/D box snoRNAs. miCLIP-sequencing RT stops and mapped reads on SNORD3/U3 and SNORD118/U8. miCLIP-sequencing data were aligned to human SNORD3/U3 (ENST00000620232) or SNORD118/U8 (ENST00000363593). The mapping information was retrieved using Samtools. Reverse transcription (RT) stops on SNORD3/U3 **(A)**, SNORD118/U8 **(B)** per 1000 mapped reads were plotted by assigning the start (+1) sites of their respective Read1 sequence. miCLIP reads mapped to SNORD3/U3 **(C)** and SNORD118/U8 **(D)** in NOP2/NSUN1 WT or C459A mutant IP samples normalized to the total mapped reads of each corresponding sample.

**Supplementary Figure S3.**
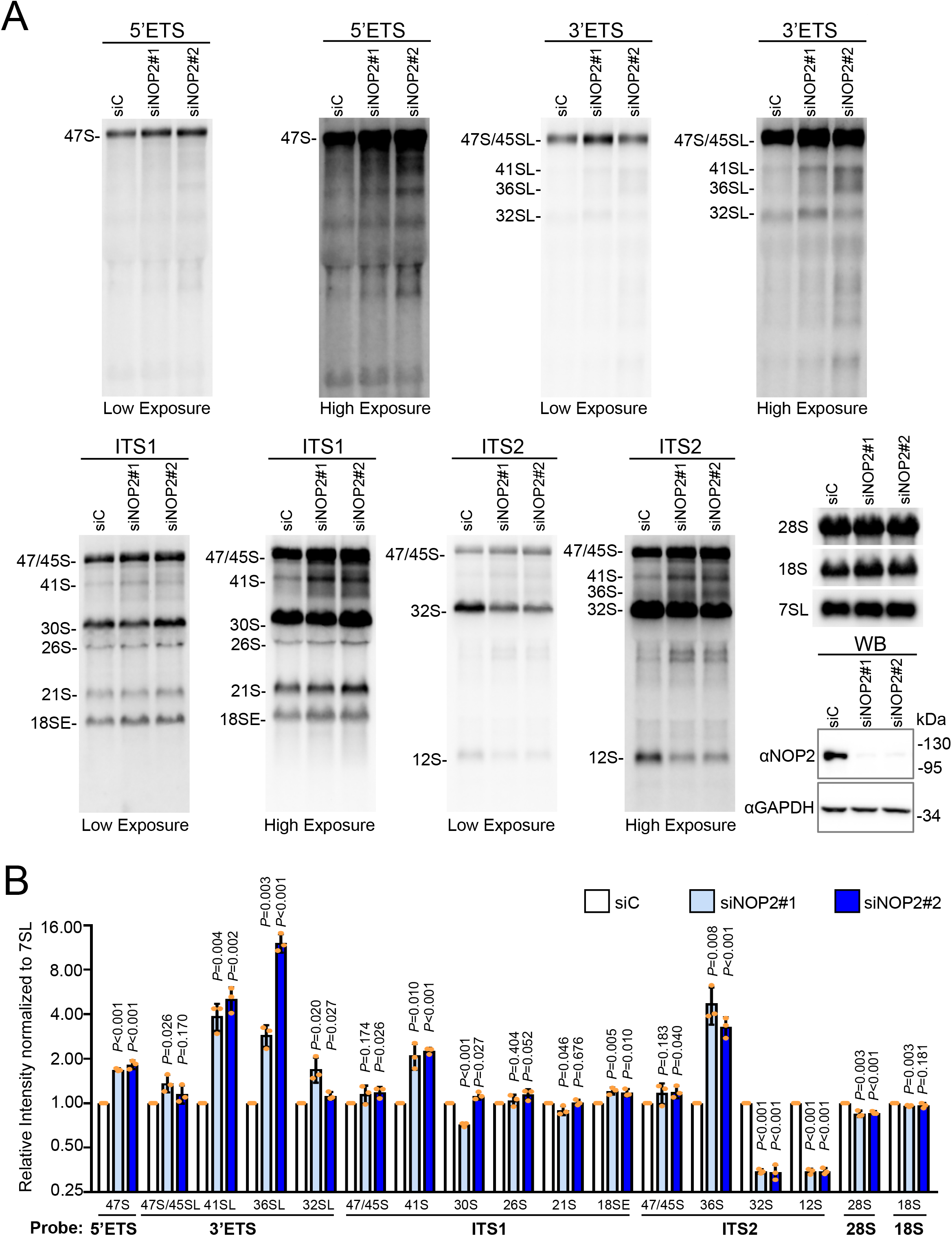
NOP2/NSUN1 is required for pre-rRNA processing. **(A)** HCT116 cells were transfected with non-targeting control (siC), NOP2 siRNA #1 or #2. After 72h, total RNA was separated on formaldehyde denaturing agarose gel and analyzed by Northern blot with 5’ETS, 3’ETS, ITS-1, ITS-2, 18S, 28S, and 7SL probes. A fraction of cells was collected to assess NOP2/NSUN1 depletion efficiency by Western blot (WB). **(B)** Densitometry quantification of each rRNA precursor from (A) normalized to 7SL RNA. The data are presented as the mean of 3 independent biological replicates ± standard deviation (SD). Statistical significance between NOP2/NSUN1 depleted samples and non-targeting siRNA control samples was calculated using a 2-tailed independent student *t*-test.

**Supplementary Figure S4.**
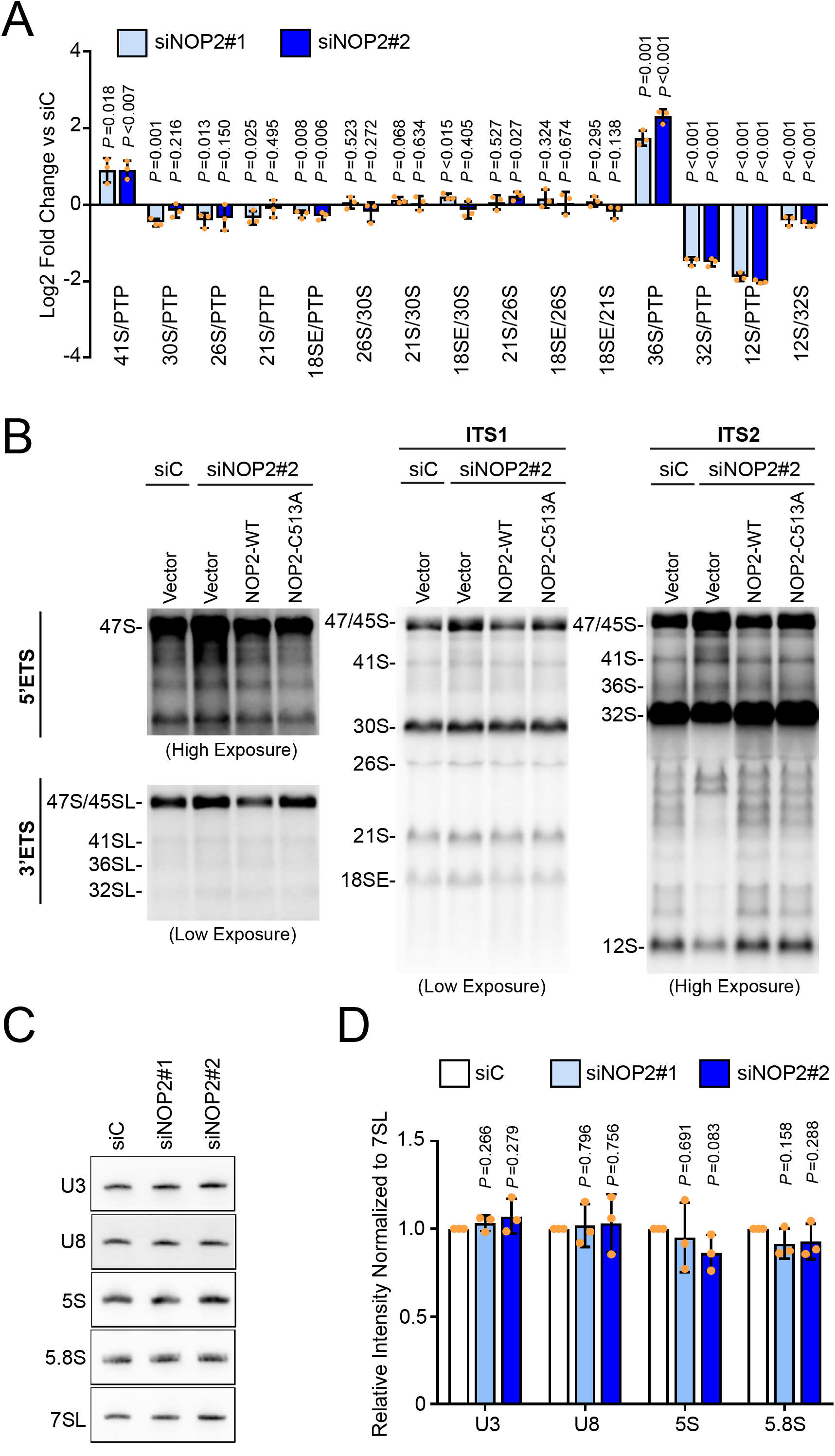
NOP2/NSUN1 depletion impairs pre-rRNA processing but does not affect SNORD3/U3 and SNORD118/U8 stability **(A)** Densitometry ratio between each two rRNA precursors from Northern blot in Figure S3A. Data are presented as the mean of 3 independent biological replicates ± standard deviation (SD). Statistical significance between NOP2/NSUN1 depleted samples and non-targeting control samples was calculated using a 2-tailed independent student *t*-test. **(B)** Different exposure of Northern blots from Figure 5B. **(C)** RNA samples from Figure S3A were separated on urea-PAGE gel and analyzed by Northern blot with SNORD118/U8, SNORD3/U3, 5S, 5.8S, and 7SL specific probes. **(D)** Densitometry quantification of SNORD118/U8, SNORD3/U3, 5S, and 5.8S signal from (B) normalized to 7SL RNA signal. Data are presented as the mean of 3 independent biological replicates ± standard deviation (SD). Statistical significance between NOP2/NSUN1 depleted samples and non-targeting control samples was calculated using a 2-tailed independent student *t*-test.

**Supplementary Figure S5.**
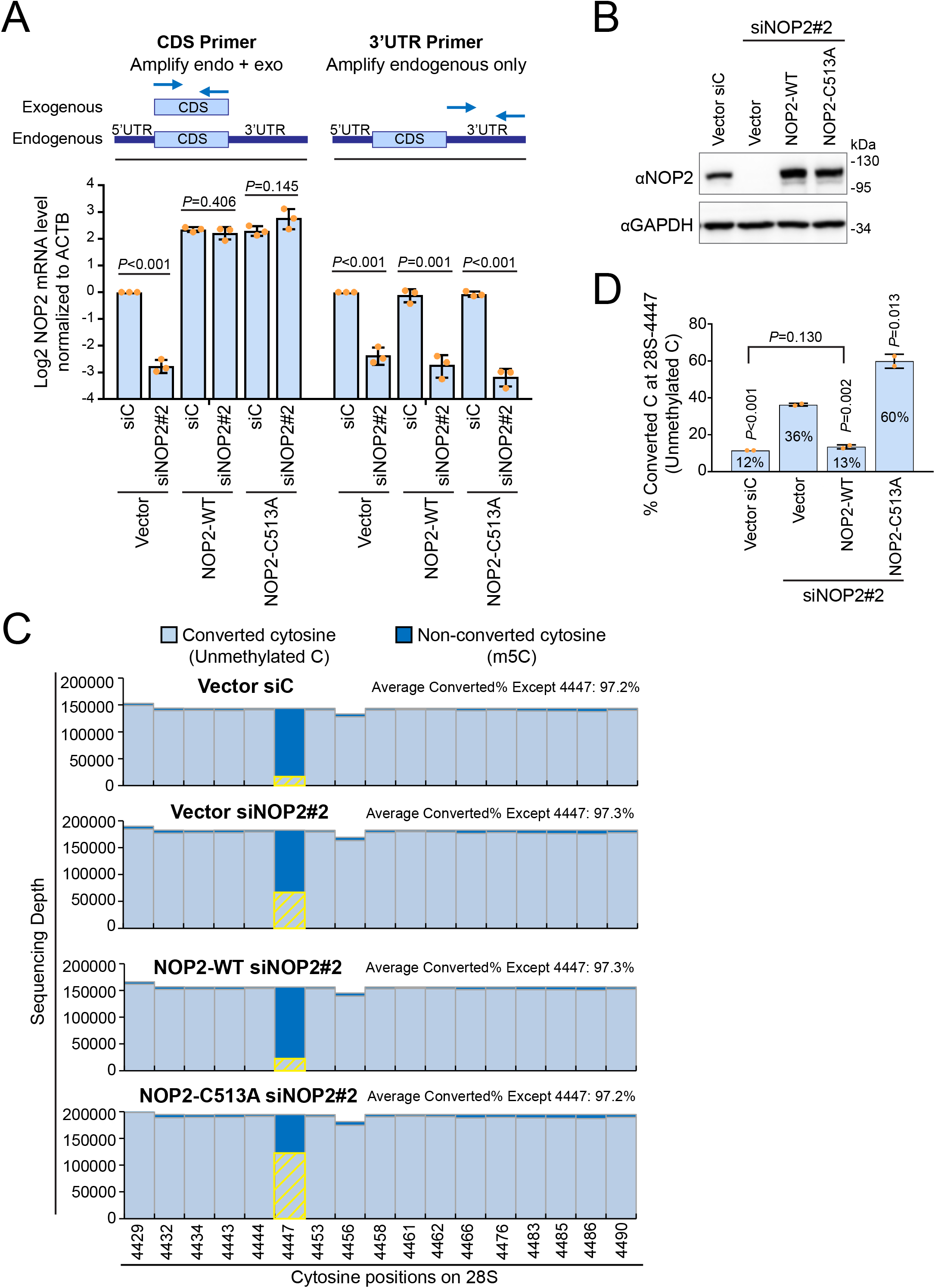
NOP2/NSUN1 cysteine 513 is required to catalyze m5C modification at position 4447 of the 28S rRNA. **(A)** HCT116 cells expressing empty vector (Vector), siRNA resistant NOP2/NSUN1 WT, or C513A mutant were transfected with non-targeting control (siC) or NOP2 siRNA #2. After 72h, total RNA was extracted and NOP2/NSUN1 depletion was assessed by RT-qPCR. Primers specific to the *NOP2* coding sequence (CDS) were used to detect both endogenous and exogenous *NOP2* and primers specific to *NOP2* 3’UTR were used to detect endogenous *NOP2* only. Expression of *ACTB* was used for normalization. RT-qPCR data are presented as the mean of 3 independent biological replicates ± standard deviation (SD). Statistical significance between NOP2 depleted samples and non-targeting siRNA control samples was calculated using a 2-tailed independent student t-test. **(B)** A fraction of cells from (A) were used to determine NOP2/NSUN1 depletion efficiency by Western blot with the indicated antibodies. **(C)** A fraction of cells from (A) were harvested for nuclear RNA extraction. Nuclear RNA was treated with bisulfite salt and sequenced on the Illumina Mi-Seq platform. Numbers of non-converted (5mC modification) and converted cytosines (unmethylated C) for each cytosine position near C4447 were plotted. Yellow marks highlight converted cytosines at position 4447. **(D)** Percentage of converted cytosine (C) at position 4447 on the 28S rRNA in each sample. Data are shown as the mean of 2 independent biological replicates ± standard deviation (SD). Statistical significance values relative to empty vector + NOP2/NSUN1 knockdown (Vector siNOP2#2) were calculated using a 2-tailed independent student t-test.

**Supplementary Figure S6.**
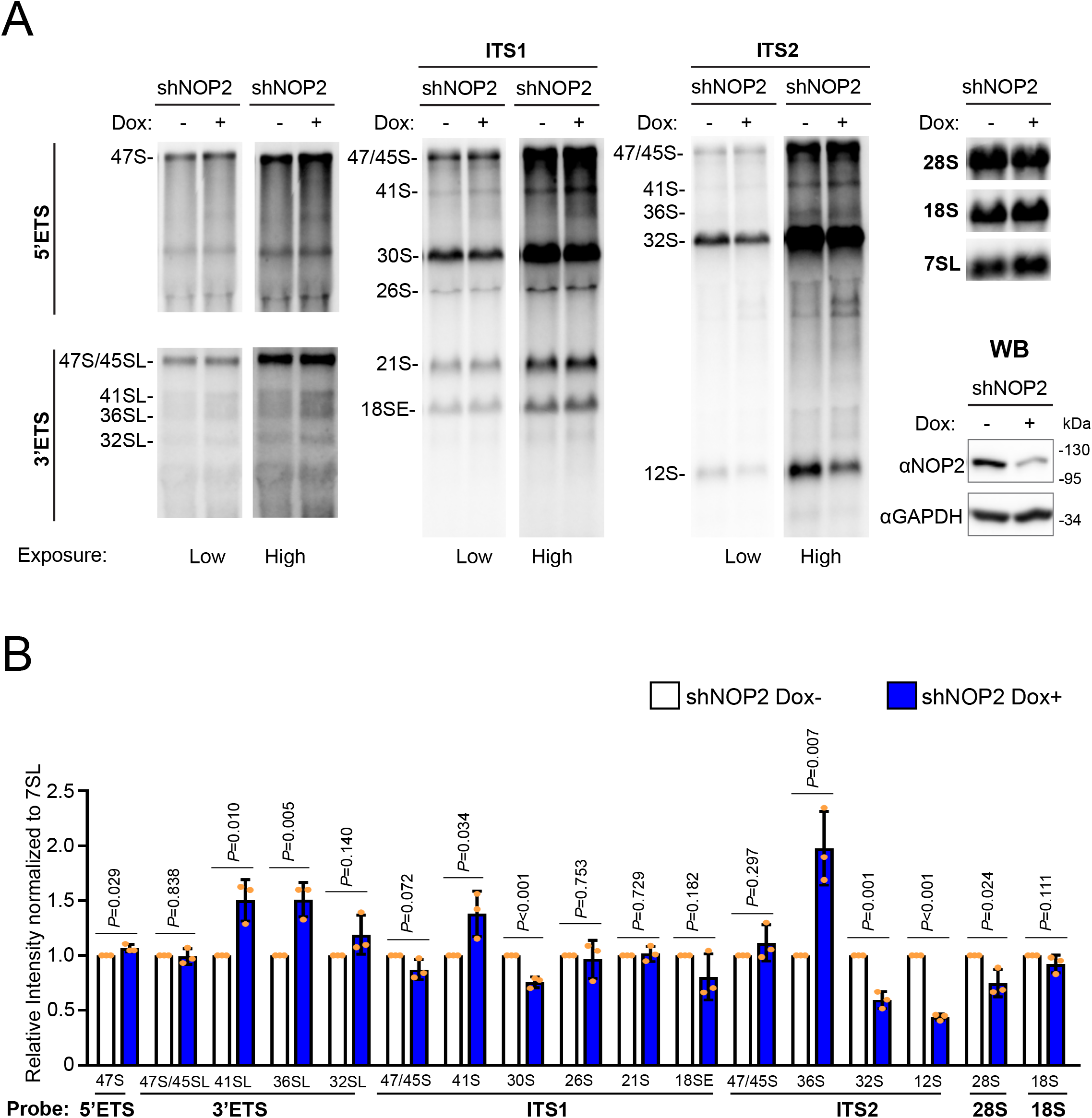
Depletion of NOP2/NSUN1 by inducible shRNA causes rRNA processing defects. **(A)** HCT116 cells expressing doxycycline-inducible NOP2 shRNA were induced with 200 ng/ml doxycycline (Dox+) for 4 days. Non-induced (Dox-) cells were used as control. Total RNA was extracted and separated on formaldehyde denaturing agarose gel and analyzed by Northern blot with 5’ETS, 3’ETS, ITS-1, ITS-2, 18S, 28S, and 7SL probes. A fraction of cells was collected to determine NOP2/NSUN1 depletion efficiency by Western blot (WB) with the indicated antibodies. **(B)** Densitometry quantification of each rRNA precursor from (A) normalized to 7SL RNA. The data are presented as the mean of 3 independent biological replicates ± standard deviation (SD). Statistical significance between NOP2 depleted samples (Dox+) and non-induced control samples (Dox-) were calculated using a 2-tailed independent student t-test.

**Supplementary Figure S7.**
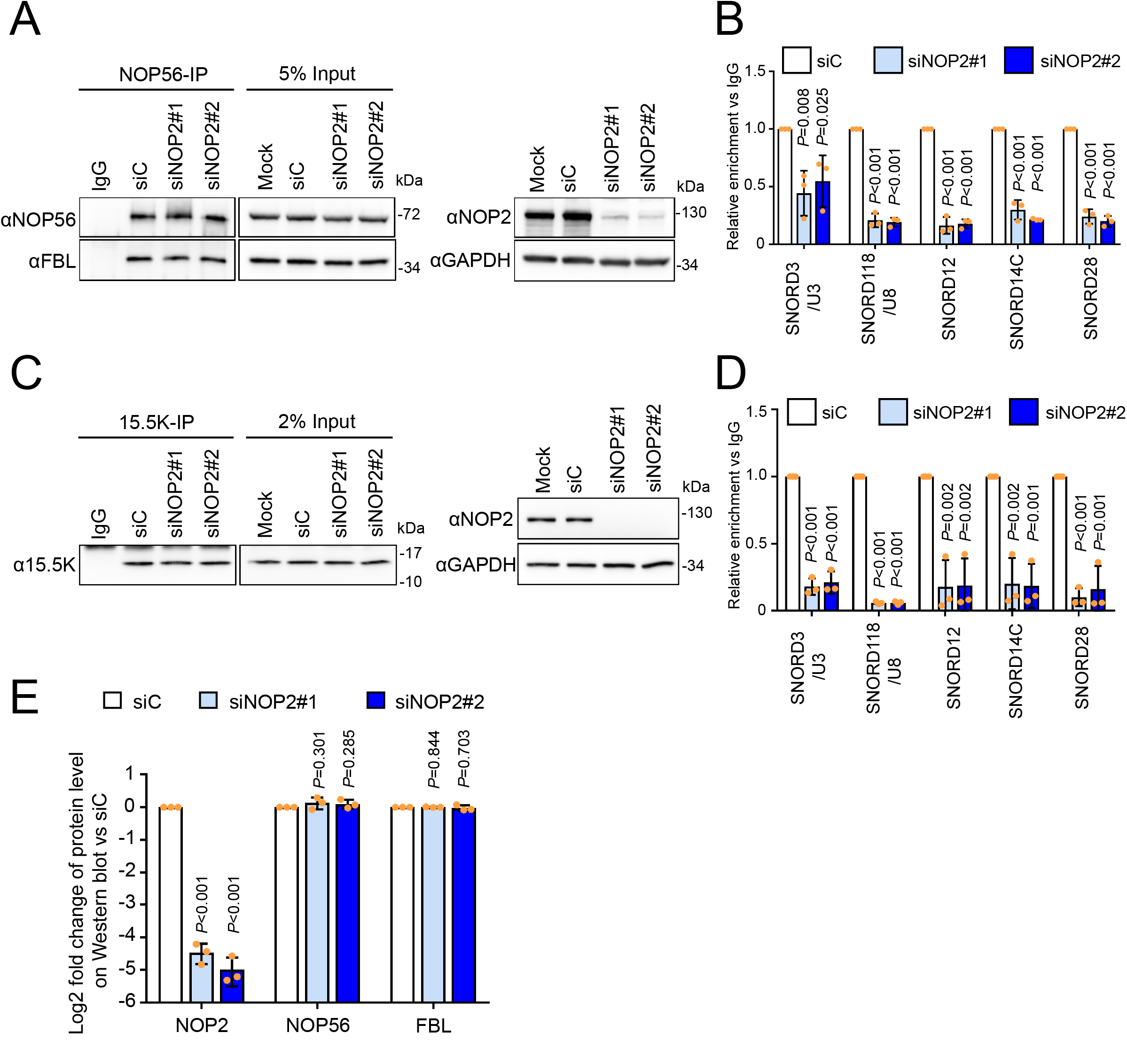
NOP2/NSUN1 is required to maintain the integrity of C/D box snoRNPs. HCT116 cells were transfected with non-targeting control (siC), NOP2 siRNA #1 or #2. 72h later, cells were lysed and immunoprecipitated (IP) with anti-NOP56 antibody **(A, B)** or 15.5K antibody **(C, D)**. A fraction of the immuno-precipitate was analyzed by Western blot to control for immunoprecipitation and knockdown efficiency **(A, C)**. The remaining immunoprecipitated fraction was processed for RNA extraction. NOP56 **(B)** or 15.5K **(D)** associated RNA was analyzed by RT-qPCR with SNORD3/U3, SNORD118/U8, SNORD12, SNORD14C, and SNORD28 specific primers. Relative enrichment over IgG control is represented. The Data is presented as the mean of 3 independent biological replicates ± standard deviation (SD). Statistical significance between NOP2/NSUN1 depleted samples and non-targeting siRNA control samples was calculated using a 2-tailed independent student *t*-test. **(E)** Densitometry quantification of each protein in input samples from (A) normalized to GAPDH. Data are presented as the mean of 3 independent biological replicates ± standard deviation (SD). Statistical significance between NOP2 depleted samples and non-targeting control samples was calculated using a 2-tailed independent student t-test.

**Supplementary Figure S8.**
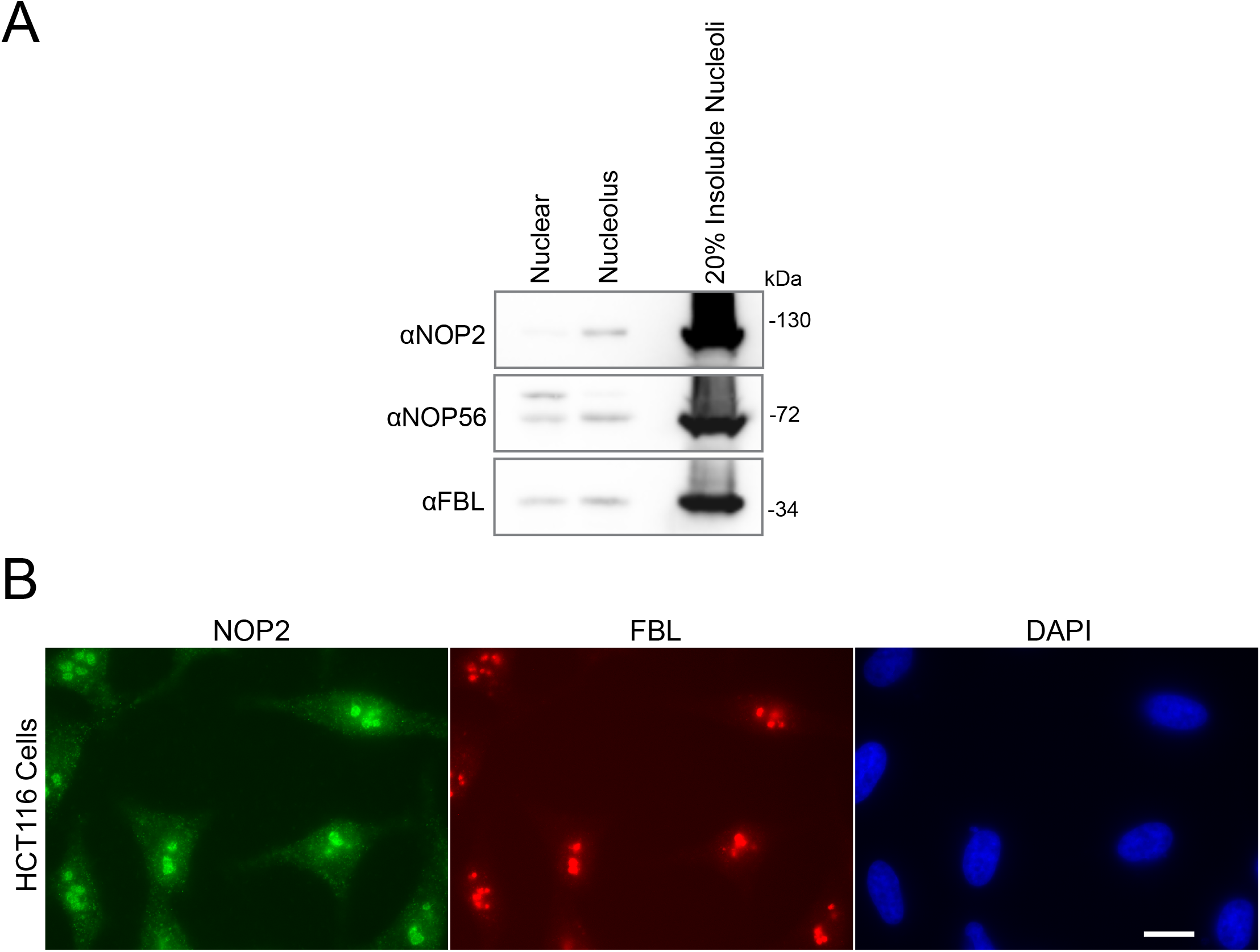
Human NOP2/NSUN1 is predominantly located in nucleoli. **(A)** Proteins in nuclear extracts, nucleolar extracts, and insoluble nucleoli pellets from HCT116 cells were analyzed by Western blot with the indicated antibodies. **(B)** HCT116 cells were fixed and immunostained with anti-NOP2/NSUN1 (green) and anti-FBL (red, nucleolar marker). DNA was visualized by staining with Hoechst 33342 (blue). Scale bar is representative of all panels: 10 μm.

**Supplementary Figure S9.**
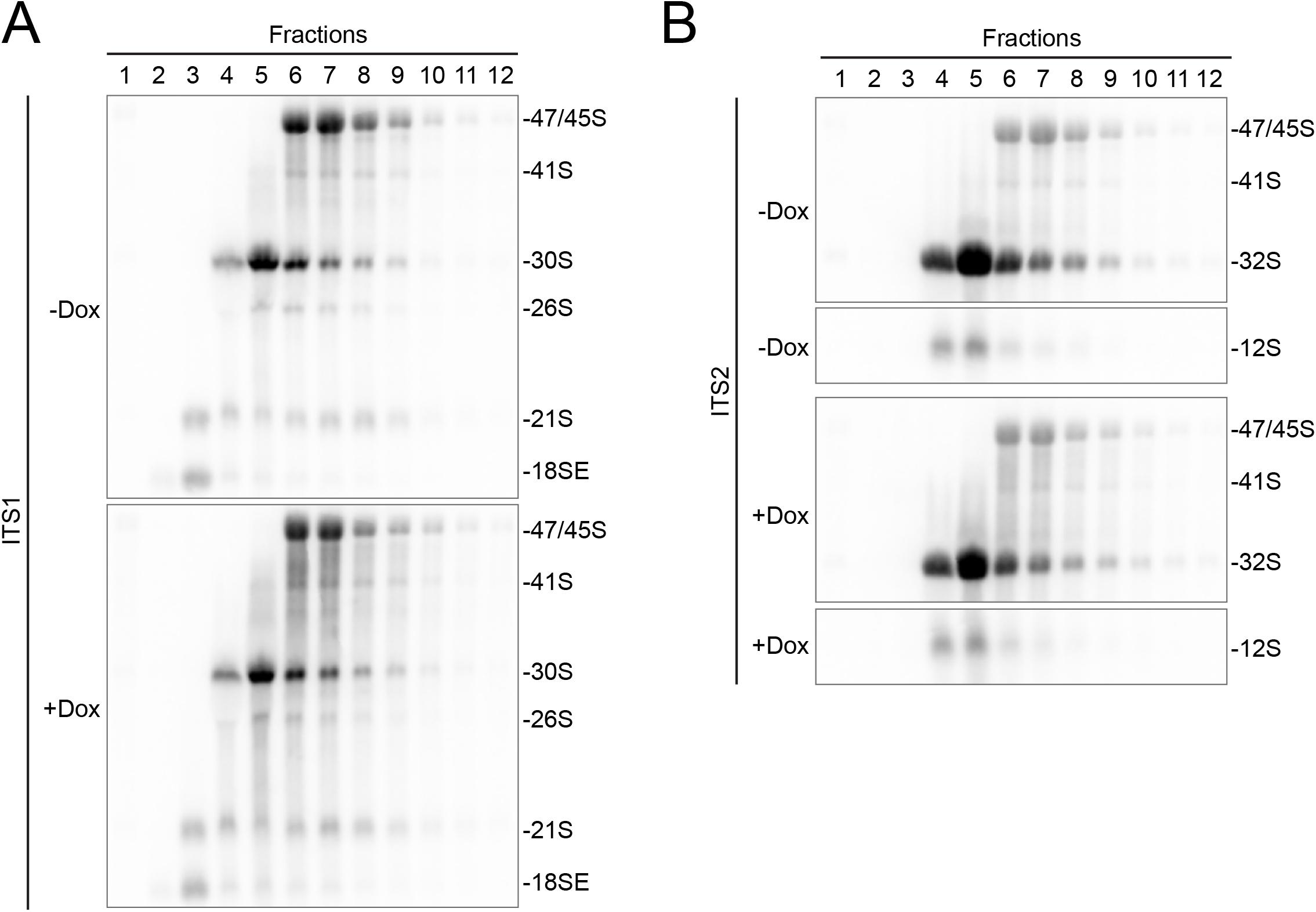
Northern blot analysis of rRNA precursors present in each fraction from the sucrose gradient fractionation from Figure 9A. RNA from fractions 1-12 was extracted, separated on formaldehyde denaturing agarose gel, and analyzed by Northern blot with ITS1 **(A)** and ITS2 **(B)** probes.

**Supplementary Figure S10.**
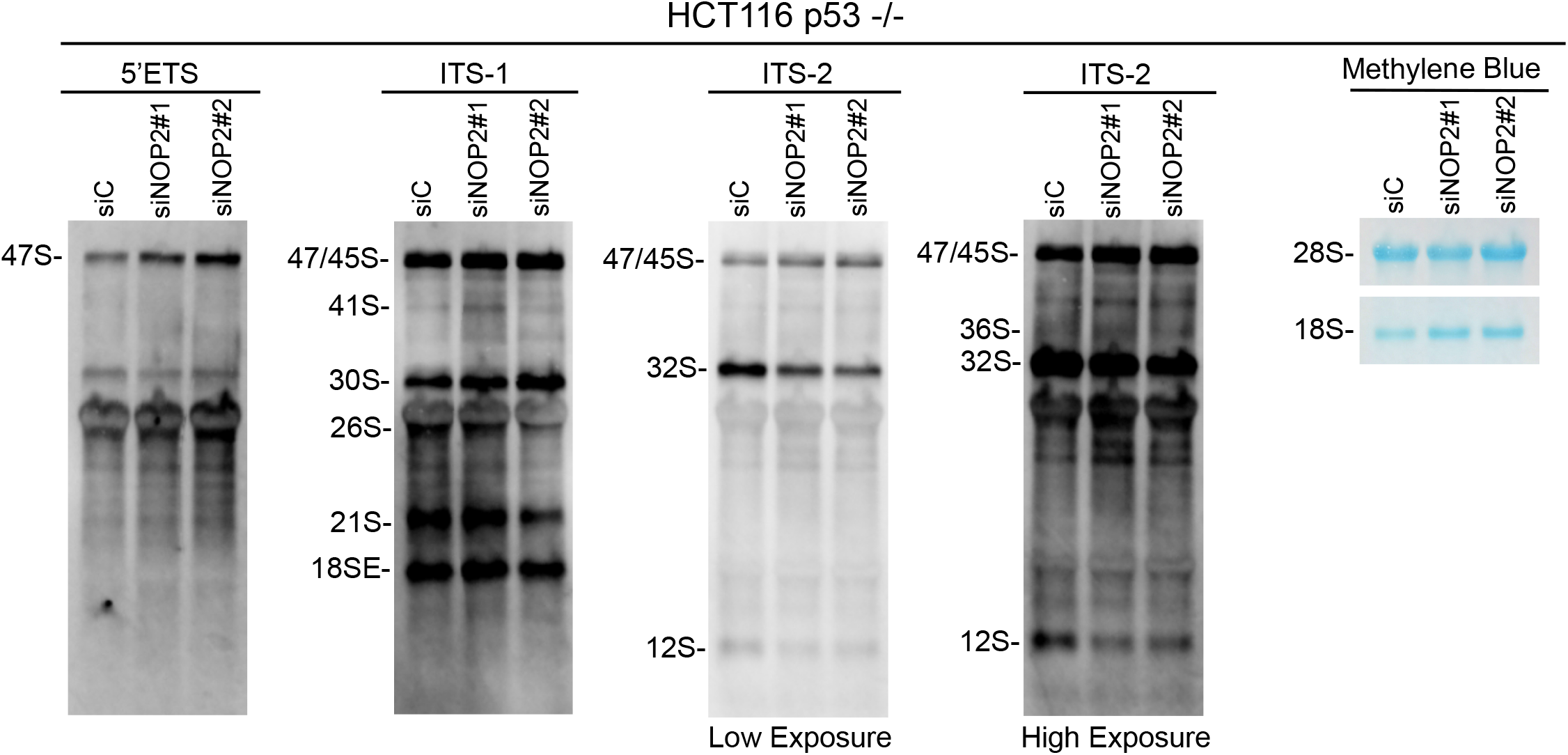
Pre-rRNA processing defects observed after NOP2/NSUN1 depletion are independent of p53 status. HCT116 p53 -/- cells were transfected with non-targeting control (siC), NOP2 siRNA #1 or #2. 72h later, total RNA was separated on formaldehyde denaturing agarose gel and analyzed by Northern blot with 5’ETS, ITS-1, and ITS-2 probes. Right panel: 28S and 18S rRNA were stained with methylene blue on the membrane.

**Supplementary Table 1.** Sequence of primers and probes used in this paper.

**Supplementary Table 2.** miCLIP sequencing Peaks with IP vs SMInput fold-change >= 2 and adjusted P value < 0.05.

